# Thalamocortical, not Intracortical, Inputs Set the Wakeful Activity Levels of Cortical Visual Inhibition

**DOI:** 10.64898/2026.10.14.759116

**Authors:** Jérémie Sibille, Kai Lun Teh, Jasper Alus Pysall, Jens Kremkow

## Abstract

Cortical neurons integrate from a wide range of synaptic inputs, locally within the cortex and distally from multiple brain sources, which altogether support the generation and transmission of spikes. Yet, the relative influence on local cortical processing between thalamocortical and intracortical inputs still remains poorly understood. Here, we used tangential insertion of high-density silicon probes to capture thalamocortical next to intracortical functional connectivity, thereby unraveling their relative contribution to local cortical dynamics. By comparing wakefulness versus light anesthesia, we revealed a marked wakeful increase of firing in specifically thalamocortical axons and putative cortical inhibitory neurons; an increase which is very homogeneous in both populations, and comparatively absent in neighboring cortical excitatory neurons. Such specific comodulation is attributable to the known preferential connectivity of thalamocortical axons toward putative cortical inhibitory neurons, which we quantified to follow a 90/10% preference. In addition, we uncovered that wakeful increases of activity levels are symmetric in thalamocortical synapses and asymmetric in intracortical synapses. Indeed, within intracortical synapses, the wakeful increases of firing mostly happen in receiving neurons, leaving pre-synaptic upstream neurons barely impacted by wakefulness. In the meantime, thalamocortical receiving excitatory neurons exhibit a marked wakeful increase of firing, unlike the rest of the excitatory cortical population. Using mathematical modelling simulating spike transmission in thalamocortical and intracortical networks, we showed that only outnumbering thalamocortical-to-intracortical inputs could mimic the observed homogeneous wakeful increases of firing in thalamocortical and cortical inhibitory neurons. Altogether, our results identify thalamocortical synaptic inputs as a key controller of the wakeful level of firing of inhibitory neurons at the first relay of visual cortical processing.

## INTRODUCTION

The cortical mantle is in essence hosting billions of synaptic connections, between millions of neurons, even in a small mouse brain. This intricate network is organized according to a certain set of rules, starting with the fact that: almost all mammalian cortices are made of six canonical layers, organized as parallel modular columns (Mountcastle, 1997; Harris and Mrsic-Flogel, 2013). Each layer receives and sends a distinct set of connections, following certain forms of patterns (Goulas et al., 2019; Puxeddu et al., 2024). In simplified terms, layer 1 receives connections from other cortical regions (Doron et al., 2020), whereas layer 2-3 forms more localized connections (Jouhanneau et al., 2018), with a preference toward similarly encoding neurons (Ko et al., 2013; Cossell et al., 2015). Layer 4 receives direct thalamic inputs, while layers 5 and 6 are hosting multiple neuron types, separable by their downstream synaptic targets, either toward the thalamus, sub-thalamic targets, or other cortical or striatal regions (Bodor et al., 2025; Moberg et al., 2022). In an oversimplified scheme, layer 5 and 6 neurons are the output of the cortical column broadcasting the local spiking to other cortical areas (layer 5) or directly to the thalamus (layer 6). Layer 6 neurons are known to be instrumental to the tonicity of primary thalamic nucleus (Dimwamwa et al., 2024, Olsen et al., 2012, Born et al., 2021) whereas they include a sub-population having mostly cortical downstream targets which are associated to attention (Zolnik et al., 2024; 2026). Consequently, the cortical mantle is heavily interconnected with the thalamus, which acts both as a sensory gateway (Usrey et al., 2015) and as an “orchestra conductor” (Llinas et al., 1998; 2001). Each sensory cortical column receives inputs from its primary thalamic nucleus, mostly in layer 4 (Bopp et al., 2017; Constantinople et al., 2013; Kremkow et al., 2018), and to a lesser extent in layer 5 and 2-3 (Harris et al., 2019), while secondary thalamic nuclei preferentially target upper cortical layers 1, 2-3 and deeper layer 5-6 (Zhang and Bruno, 2019). Functionally, paired recordings *in vivo* unravelled that excitatory connections exhibit some preference toward narrow-waveform neurons (putative fast-spiking interneurons), as observed in thalamocortical connections (Bruno and Sakmann, 2006; Swadlow and Gusev, 2002), and layer 2-3 intracortical connections (Jouhanneau et al., 2023).

Together, these intermingled neuronal networks form the substrate of all cortical computations, from local microcircuits to global brain states. Yet how these neuronal circuits integrate sensory cues and generate plans and consequent actions remains largely unresolved. Therefore, to tackle these complex questions, anesthesia provides a simplifying means to unravel certain aspects underlying wakeful brain processing. General anesthesia is characterized by a loss of consciousness thought to depend on cortical (Koch et al., 2016; Brown et al., 2011; Hudetz and Mashour, 2016), and thalamic (Whyte et al., 2024) activity. Yet, different sensory cortical areas continue to respond to sensory inputs under anesthesia (Haider et al., 2013; Filipchuk et al., 2022; Bruno et al., 2006); thereby raising the question of which specific changes impact brain consciousness. Proposed explanations include alterations in single neuronal physiology (Suzuki et al., 2020), impaired assembly recruitment (Filipchuck et al., 2022), imbalance in excitation-inhibition balance (Haider et al., 2013), disrupted communications between brain areas (Alkire et al., 2000), or loss of noradrenalinergic tonus (Constantinople et al., 2011). General anesthesia is also associated with slow, synchronized cortical activity patterns (Hagihira, 2015; Lee et al., 2013; Steriade et al., 1993), peaking in the alpha band (Cartailler et al., 2019), which is more prominent in layer 5 neurons (Bharioke et al., 2022).

In the last decade, high-density silicon probes (Jun et al., 2016; Steinmetz et al., 2019; Ye et al., 2025) have broadened the possibility to record multiple neurons from multiple brain areas simultaneously (Siegle et al., 2021; Chen et al., 2024; Durand et al., 2023). These probes can also capture non-somatic spiking activity, in particular axons across vertebrate brain (Ye et al., 2025), back propagating dendritic action potentials in the mice cortex (Jia et al., 2019), axonal arbors, in the midbrain (Sibille et al., 2022; Gehr et al., 2023), and in the cortex (Sibille et al., 2024). Notably, even earlier-generation silicon probes had already captured some non-somatic spiking activity (Sun et al., 2021; Someck et al., 2023), see review (Sibille et al., 2026). The possibility to measure axonal arbors has opened the door to more systematic measures of synaptic connections *in vivo*, leveraging finer monitoring of the brain physiology, as done here in wakeful and anesthetized brain states.

Collectively, these studies have separately tackled several aspects underlying wakeful physiology, by characterizing the processes occurring under anesthesia: but they all lacked the observation(s) of putative concomitant changes of cortical versus thalamocortical functional connectivity, which the present study proposes to address. Expanding from the capability to capture functional thalamocortical connectivity (Sibille et al., 2024), we now look into neighboring intracortical connectivity, asking how their balanced integration supports wakeful cortical physiology *in vivo*.

## Results

### Wakefulness increases thalamocortical axons and narrow-waveform neuronal activity

To collectively observe two different neighboring synaptic contacts at the first relay of cortical visual processing, we used high-density silicon probes to monitor converging spiking activity in the primary visual cortex. Using previously established methods (Sibille et al., 2024), we inserted Neuropixels probes tangentially in the primary visual cortex of head-fixed mice which were passively exposed to classical visual stimuli on the screen (Fig. 1a, see methods). We extracted thalamocortical axons (TCA, cf methods for controls), broad waveform neurons (BWN, putative excitatory), and narrow-waveform neurons (NWN, Fig. 1b, see materials and methods and Sup. Fig. 1), the latter being an approximation for inhibitory fast spiking neurons (Swadlow and Gusev, 2002, Dimwamwa et al., 2024, Valero et al., 2025). The tangential insertion goes through the curved cortical mantle, thereby capturing TCA, NWN, and BWN in layer 2-3, layer 4, and 5 (Sup. Fig. 2). Neighboring TCA, NWN, and BWN all exhibit characteristic features of the visual pathway, such as overlapping receptive fields (Sup. Fig. 2f, see rev. Kremkow and Alonso, 2018). After recording the activity within the cortex for an hour while, the isoflurane mask (1%) was gradually approached to the head of the animal to smoothly lead the animal into a stable anesthetized state. The anesthetized state was further recorded for an hour, re-exposing the same visual stimuli, to compare the firing rates in both wakeful versus anesthetized conditions. These comparisons unraveled wakeful increases of firing in specifically TCA and NWN, leaving BWN firing changes with a significant but small effect size (Fig. 1c-d-f, Sup. Fig. 3b, avg. rate. awake versus anesthetized (aw./ane.) TCA = 3.02 ± 0.08/0.86 ± 0.04 Hz, n = 545, NWN = 3.35 ± 0.08/1.32 ± 0.05 Hz, n = 1272, BWN = 1.38 ± 0.03/0.93 ± 0.018 Hz, n = 3219, p aw./ane. in resp. TCA/NWN/BWN = 6.9*10^-89/1.9*10^-167/2.07*10^-62, Cliff’s δ = 0.79/0.59/0.15, pooled from n = 11 mice, cf. methods). Associated with these firing changes, other firing properties were scrutinized to report more classical changes observed under anesthesia, looking into firing burstiness and coherence (Sup. Fig. 3-4). It is to be noticed that in only certain TCA, clear bursty firing patterns appear during anesthesia, indicating the presence of pairs of spikes in the alpha band (Sup. Fig. 3 c) as previously observed in the anesthetized dLGN from cats (Reinagel et al., 1999). It was puzzling to observe that the BWN activity under anesthesia is little impacted, neither by the loss of thalamic inputs, nor by the decrease of inhibitory activity (Fig. 1c-d-f, Cliff’s δ = 0.15 (small)). This absence of medium to large effect size can be attributed to the compensating firing supported by their higher synchrony in anesthetized state, as captured by their higher coefficients of variation (Sup. Fig. 4d-e, from Bharioke et al., 2022). For the rest of this study, we quantified the change in firing rate (or other properties), using a modulation index 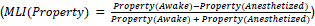. When looking at the MLI of firing, we first quantified their changes based on comparison to equivalent distributions of zeros (Fig. 1c, in resp. TCA/NWN/BWN: MLI avg. = 0.576 ± 0.01/0.474 ± 0.009/0.123 ± 0.007, n = 545/1272/3219, one-sample Wilcoxon signed-rank test, p = 3.01*10^-151/6.7*10^-220/1.6*10^-28, Cliff’s δ = 0.95/0.54/0.21). To highlight the ubiquitous wakeful increase of firing, we reported the MLI of firing in each individual mouse (Fig. 1f, from n = 11 mice). Last, comparing MLI of firing between populations confirmed that TCA showed the strongest MLI, compared to NWN and to BWN (p TCA to NWN/TCA to BWN/BWN to NWN = 1.9*10^-11/ 4.5*10^-132/ 1.08*10^-161, δ = 0.19/0.65/0.51, Mann Whitney U test). Together, these results led to: first a strong impression of co-modulations between populations of TCA and NWN (MLI δ is small: 0.19), and second that the NWN population activity is homogeneously increased by wakefulness.

**Figure 1:**
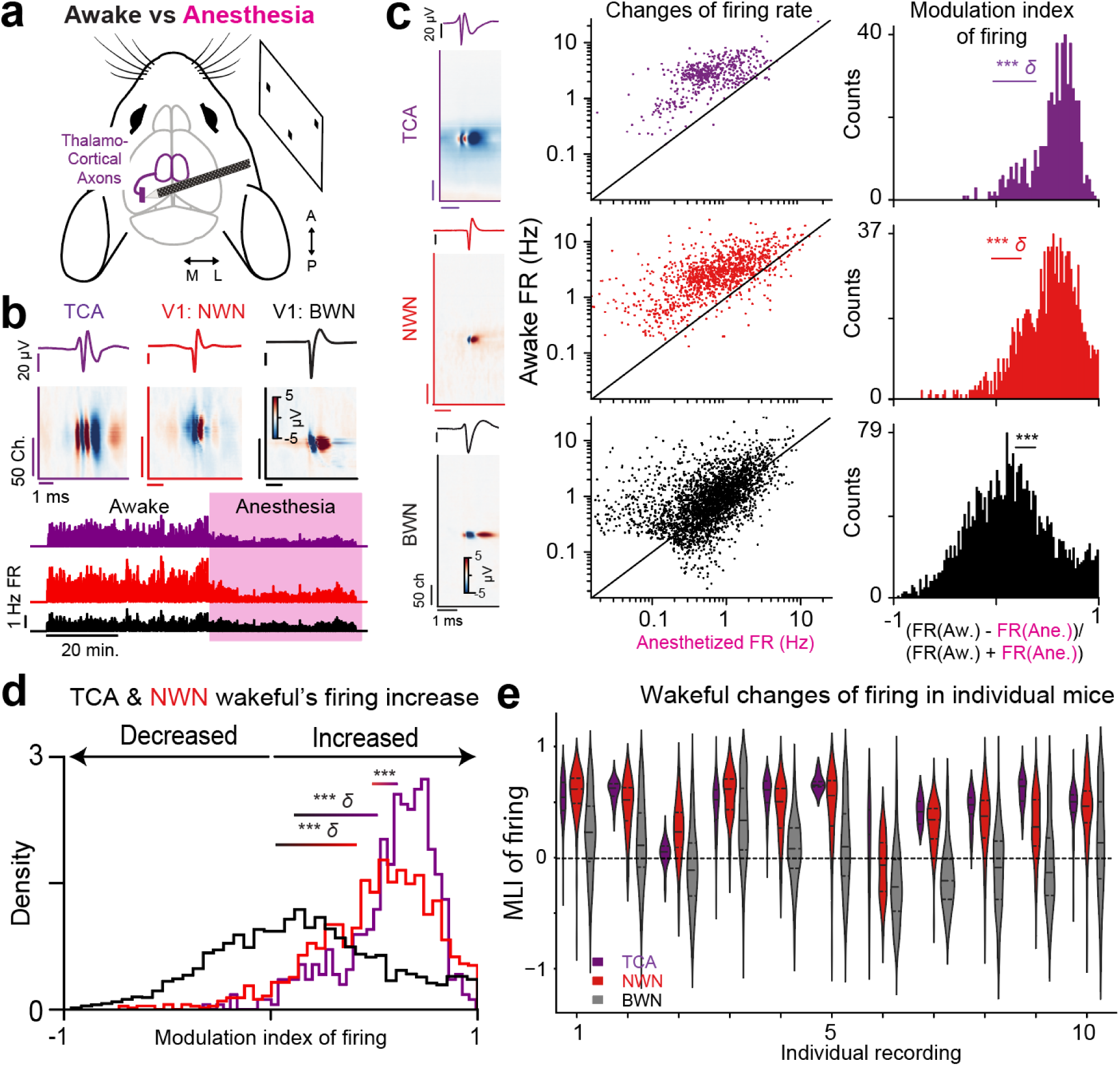
Wakefulness is associated with a specific increase of firing of both thalamocortical axons and narrow-waveform neurons in visual cortices *in vivo*. **a**, Schematic of tangential insertion with Neuropixels probes. **b**, Three types of waveforms are extracted: thalamocortical axons (TCA, purple), with their typical asymmetric double peaks; putative inhibitory narrow-waveform neurons (NWN, red), and putative excitatory broad waveforms neurons (BWN, black). All neuronal waveforms are represented by their multiple channel waveforms illustrating their electrophysiological signature on all channels of the probe, over 5 ms time, on a +-5 µV scale (top row). Illustration of the timely change of population averaged activity in a single recording as isoflurane is applied (pink, bottom) **c**, Illustration of the wakeful-induced changes of firing (log-scale axis, middle column) and the corresponding modulation indexes (MLI) of firing (right column). Statistical comparisons (indicated by a horizontal line) are done versus distributions of zero with similar size (one sample Wilcoxon signed-rank, Cliff delta (δ) indicate middle to large effect size) for TCA (top row), NWN (middle row) and BWN (bottom row). **d**, Normalized histogram of the overlapping MLI of firing of TCA, NWN and BWN populations highlighting the similar increase of firing in both TCA and NWN (n = 545 TCA/ 1272 NWN/ 3219 BWN, n = 11 mice, comparisons are done between populations using Mann-Whitney-U-test). **e**, Illustration in each mouse of the consistent wakeful increase of firing.

**Figure 2:**
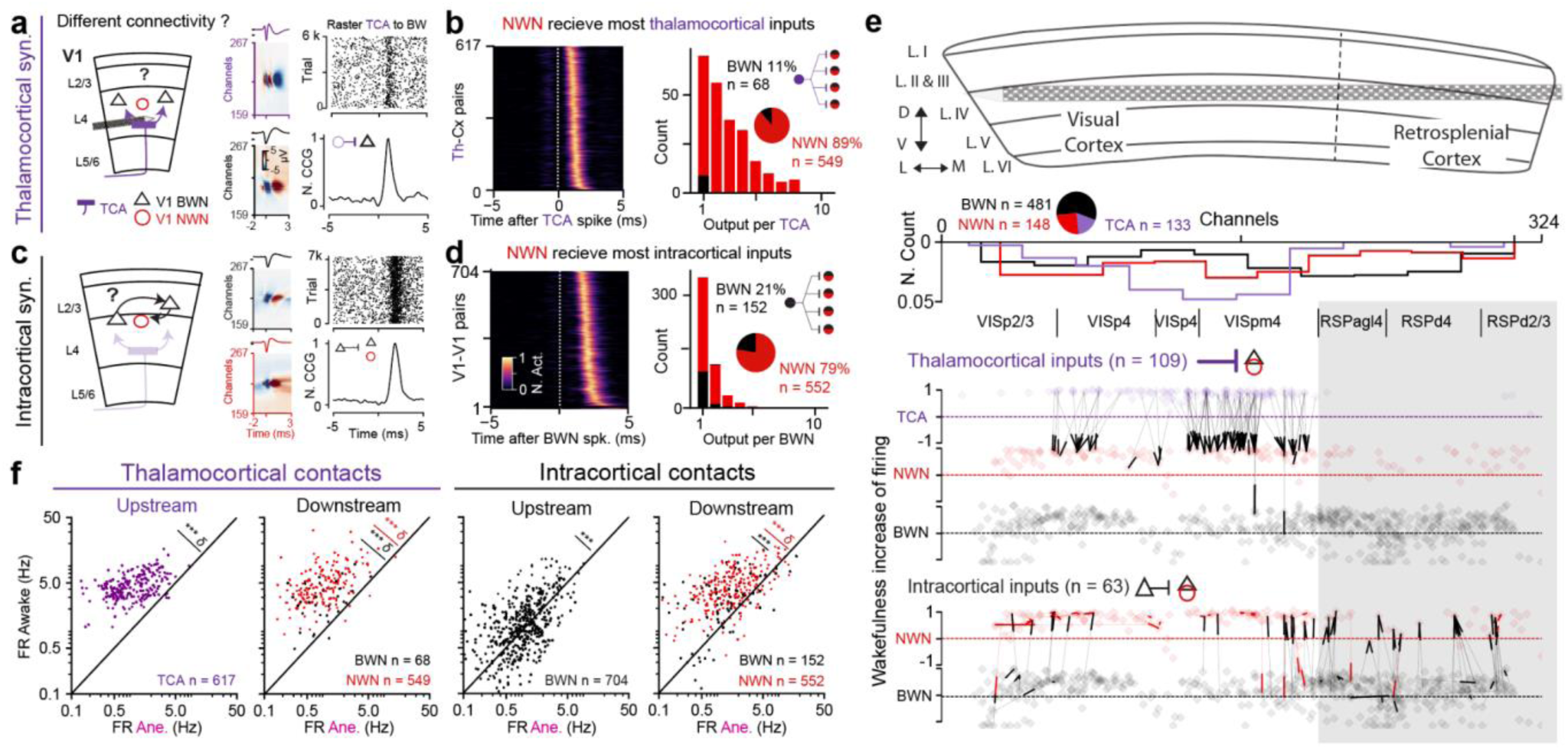
Wakefulness differently modulates upstream vs downstream neuronal firing at thalamocortical and intracortical functional synaptic contacts. **a**, Schematic of thalamocortical excitatory synaptic connections (left). Rasters between TCA and a cortical neuron revealed synchronized firing (top right), associated with a prominent peak in the corresponding cross-correlogram (CCG, bottom right). **b**, Overlapping profiles of the normalized CCGs from all detected mono-synaptic pairs between TCA and cortical neurons (left), which revealed a connectivity from TCA to mostly NWN (89% of the 617 observed pairs). In addition, thalamocortical connectivity is broadly divergent, as exemplified by the numerous post-synaptic partners (right). **c**, Same as in a, for intracortical excitatory synaptic connections, schematic (left), rasters (top right), and CCG (bottom right). **d**, Same as in b: Overlapping normalized CCG profiles (left), associated to a specific connectivity between BWN to NWN (79% of the 704 observed pairs), which is less divergent (right). **e**, Overview in a single recording of the physical location of the different observed monosynaptic contacts. Schematic of the probe insertions (top), aligned to the normalized spatial density of TCA, NWN and BWN, above the layer reconstruction (middle). Excitatory monosynaptic pairs are illustrated by a thin line with thickened endings illustrating its directions. TCA inputs (middle), aligned to BWN intracortical synaptic contacts (bottom). The y-axis shows each individual wakeful modulation index of firing. **f**, Illustration of the wakeful-increases of firing rates within thalamocortical (left) and intracortical (right) functional synaptic contacts, plotted on log-scale axis (n = 11 mice, Wilcoxon signed-rank test).

**Figure 3:**
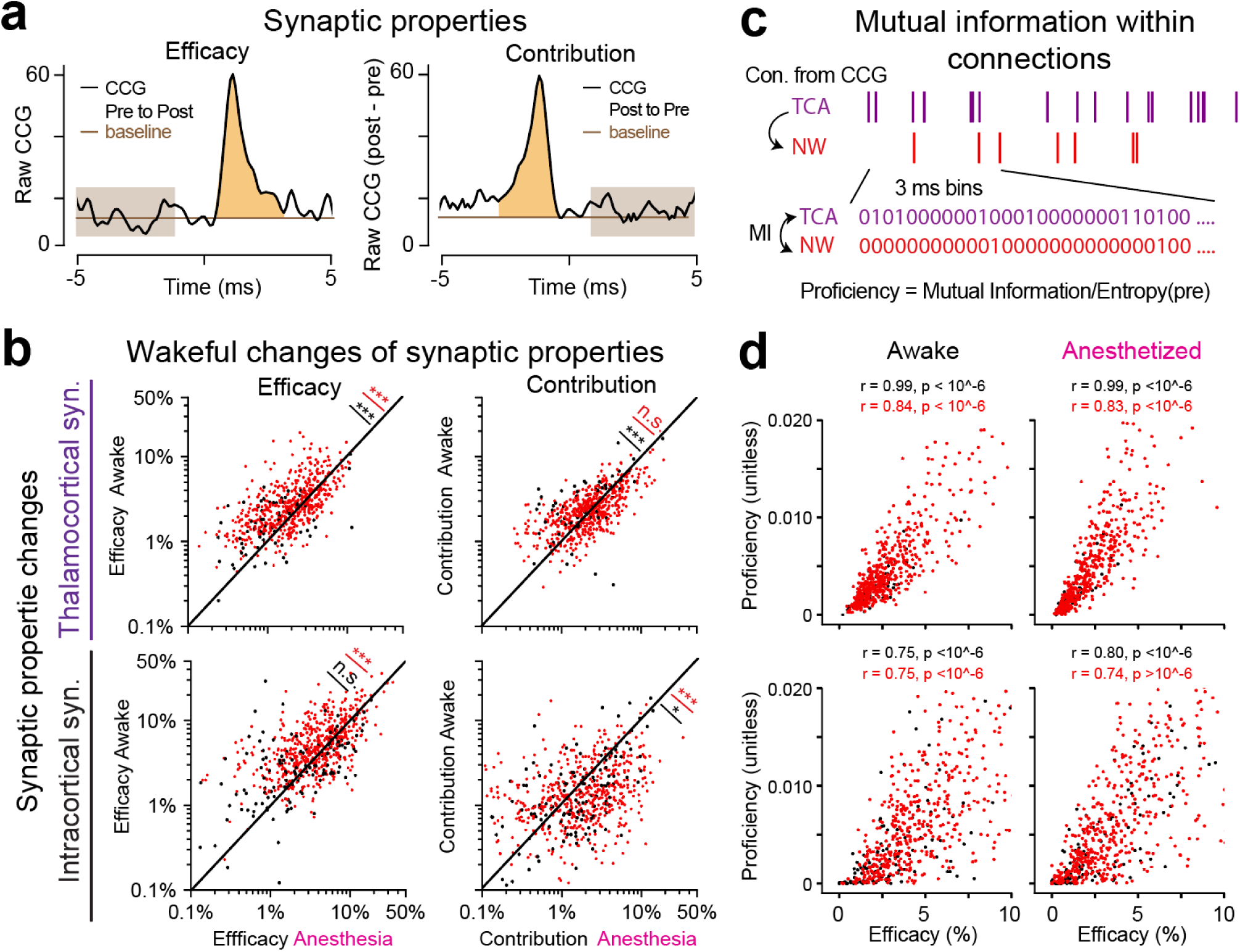
Wakeful changes of synaptic properties in thalamocortical and intracortical functional synaptic contacts. **a**, Schematic of efficacy and contribution evaluations: efficacies are estimated from the baseline-corrected pre-post CCGs (left) and contributions are estimated from the post-pre CCG (right) from the same neuronal pair. **b**, Wakeful changes of efficacies (left column) and contributions (right column) in thalamocortical (top row) and intracortical (bottom row) functional synaptic contacts. Thalamocortical synapses exhibit moderate changes of their efficacy (top left) and contribution (top right). Intracortical synapses exhibit a small increase of efficacy (bottom left), and a decrease of contribution (bottom right, Wilcoxon signed rank tests). **c**, Schematic illustration of the mutual information (MI) estimates within a functional synaptic pair. **d**, Wakeful changes of Normalized MI (Proficiency) in thalamocortical synapses (top row, toward resp. NWN n = 549, BWN n = 68 synapses) and intracortical synapses (bottom row, NWN, n = 152, BWN n = 552 synapses) in awake conditions (left column) and anesthetized (right column, n = 11 mice).

**Figure 4:**
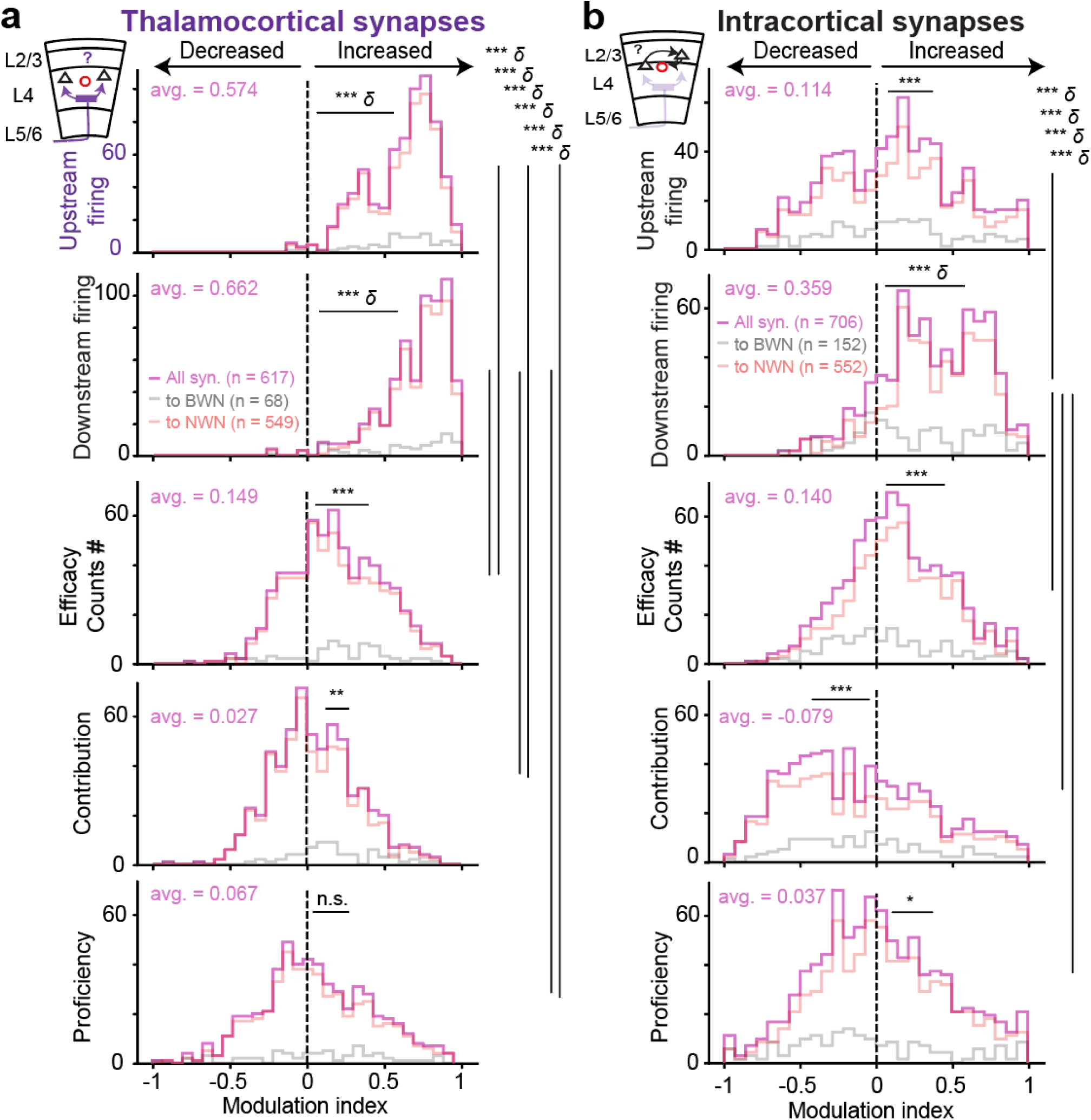
Wakefulness increase of firing is symmetric at thalamocortical synapses and only occurs in downstream intracortical neurons. **a**, Aligned wakeful modulation indexes in thalamocortical functional synaptic connections, comparing upstream firing (top), downstream firing (middle top), efficacy (middle), contribution (middle bottom), and mutual information (MI, bottom). Each plot shows the MLI for all connections (pink), for connections toward BWN (shaded gray) or NWN (shaded red). Within each plot, the above statistics correspond to MLI comparison from all connections versus zeros (one-sample Wilcoxon signed-rank, n = 617 thalamocortical connections, n = 11 mice). Between properties, statistical tests are exposed on the right side only when showing middle to large effect size (Wilcoxon signed-rank). **b**, Similar plotting for intracortical connections (n = 704 intracortical connections, n = 11 mice).

Altogether this suggests that wakefulness is associated with particular enhancement of firing levels in both TCA and NWN, when BWN population physiology seems little affected by anesthesia. Consequently, this led us to wonder whether these changes are mediated by changes within the underlying synaptic contacts. In particular, whether there are differential changes of balance between the converging thalamocortical and intracortical synaptic inputs, and how they influence cortical visual processing.

### Wakefulness-increases of firing is symmetric in thalamocortical and asymmetric in intracortical synaptic contacts

Simultaneous recording of TCA, NWN, and BWN in neighboring positions within the cortex permits to extract functional synaptic connection(s) by way of screening for prominent peaks at synaptic delays in their respective spiking crosscorrelogram (CCG, Fig. 2a-b-c-d, methods, Sup. Fig. 5). In doing so, we extracted two types of neighboring excitatory synaptic contacts: thalamocortical (Fig. 2a-b, n = 617 contacts) and intracortical functional synaptic contacts (Fig. 2c-d, n = 704 contacts, from n = 11 mice). Confirming previously published results, we observed that thalamocortical inputs connect mostly to NWN (Bruno et al., 2006; Bereshpolova et al., 2020). Given our extraction, NWN represents 28% of all cortical neurons (Sup. Fig. 1), and they receive 89% of the detected thalamocortical functional connections (Fig. 2b). Furthermore, thalamocortical functional contacts exhibit a broad diverging (Fig. 1b) and converging (Sup. Fig. 5a) connectivity scheme, similar to previous reports (Swadlow and Gusev, 2002). On the other hand, intracortical functional contacts also exhibit a preferential connectivity toward NWN, as NWN receives 79% of the intracortical synaptic contacts we detected (Fig. 2d), in line with previous publication in the somatosensory cortex (Jouhanneau et al., 2018). The extended converging/diverging connectivity scheme observed in thalamocortical contacts (Sup. Fig. 5) is not present in neighboring intracortical inputs (Fig. 2d), or any of the other types of CCG peaks we extracted (Common inputs, inhibitory inputs, see methods, Sup. Fig. 5c-d). Both types of excitatory functional synaptic contacts can be captured within a single tangential recording, but not necessarily on the same neurons (Fig. 2e). This suggests a tendency of our approach to capture thalamocortical contacts within layer 4 and deeper layers, meanwhile intracortical contacts are captured within layer 2-3; which becomes more obvious when looking into other examples (Sup. Fig. 6a-b).

**Figure 5:**
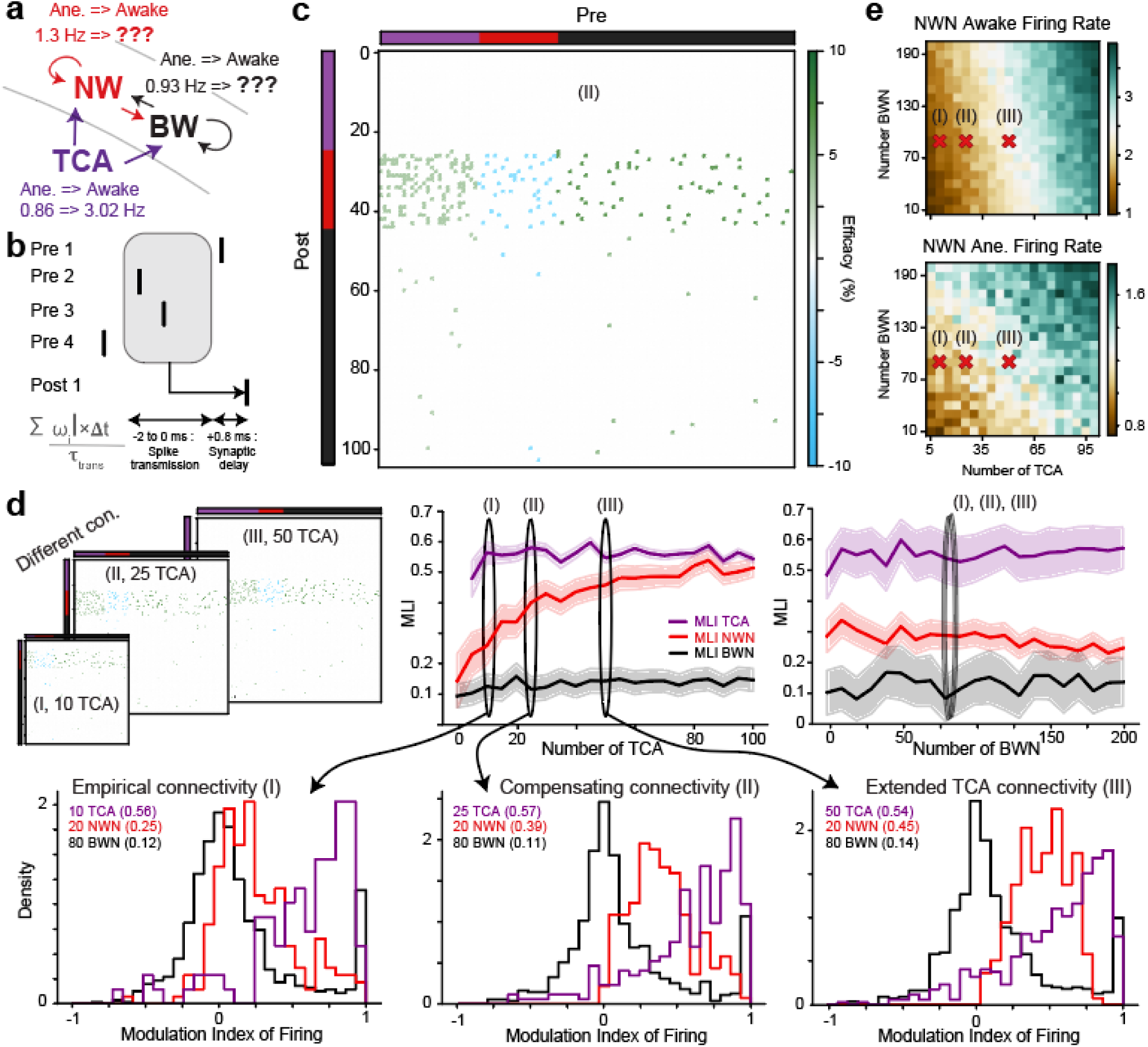
Simulating the specific NWN wakeful increase of firing in thalamo– and intracortical networks required outnumbering TCA inputs vs BWN inputs onto NWN. **a**, Schematic of the model construction. The three populations of neurons are interconnected with a fixed connectivity matrix. **b**, Schematic of spike transmission in excitatory synapses, estimated with a shifting time window of 0.8 to 2.8 ms synaptic delay, given a probability of spikes transmission from the synaptic efficacy. **c**, One connectivity matrix (“compensating connectivity”) in wakeful condition. **d**, Three connectivity scenarios (top left, I: 10 TCA, II: 25 TCA, III: 50 TCA), overlapping MLI of firing for different TCA numbers (top middle), and wakeful MLI of firing for different BWN numbers (top right). Corresponding MLI of firing in the TCA, NWN, and BWN population in the empirical connectivity matrix (bottom left, I), the compensating connectivity matrix (bottom middle), and the extended TCA connectivity matrix (bottom right) **e**, Corresponding phase diagram of NWN firing values when varying both the number of TCA (x axis) and BWN (y axis), in wakeful (top) and anesthetized (bottom) conditions.

Using these neighboring functional connected neuronal pairs, we next investigated how the changes of firing in the populations of TCA, NWN and BWN correspond to changes of firing upstream and downstream of these two excitatory synaptic contacts. We confirmed within thalamocortical contacts the wakeful increase of firing of TCA and their related downstream synaptic NWN (Fig. 2f, avg. rate TCA aw./ane. = 4.9 ± 0.08/1.4 ± 0.04 Hz, p = 1.7*10^-102, δ = 0.91, n = 617 synaptic contacts; avg. rate of downstream NWN aw./ane = 6.7 ± 0.23/1.2 ± 0.05 Hz, p = 3.2*10^-91, δ = 0.92, n = 549, n = 11 mice). In addition, we were surprised to observe in thalamocortical contacts that downstream BWN also exhibited a significant increase of firing during wakefulness, unlike the rest of the BWN population (avg. rate connected BWN awake/ane. = 3.4 ± 0.24/0.9 ± 0.14 Hz, p = 1.02*10^-12, δ = 0.8, n = 68, n mice = 11). In contrast, in intracortical functional synaptic contacts, presynaptic/upstream BWN firings exhibited similar firing levels in wakeful versus anesthetized conditions; with significant differences but negligible effect size (avg. rate upstream BWN aw./ane. = 1.6 ± 0.06/1.2 ± 0.04, n = 704, n mice = 11, p = 1.7*10^-12, δ = 0.13 is negligible, Romano et al., 2006). When asymmetrically, receiving BWN and NWN exhibit significant firing changes, with effect sizes which are medium to large (Fig. 2f, avg. rate downstream BWN aw./ane. = 3.48 ± 0.2/2.41 ± 0.24, n = 152, p = 1.1*10^-9, δ = 0.33 (medium); avg. rate downstream NWN aw./ane. = 5.15 ± 0.15/2.48 ± 0.11, p = 6.03*10^-73, δ = 0.56 (large), n = 552, n = 11 mice). This discrepancy between intracortical synaptic upstream and downstream neuronal wakeful firing changes argues that intracortical downstream neurons are under the influence of undetected thalamocortical inputs. Equally, the homogeneous wakeful increase of firing in NWN’s population (Fig. 1) suggests that all NWN we recorded from are under some TCA’s influence.

### Wakefulness induces small changes of synaptic properties

Given the specificity of wakeful firing changes in these different synaptic contacts, it leads us to wonder whether synaptic property changes could produce these different effects (Fig. 3a-b). To do so, we distinguished synaptic efficacy from synaptic contribution, which are respectively estimated from the pre-post versus the post-pre CCGs in the same pair of neurons (Fig. 3a, see methods). These two quantities reflect the probability of spike transmission being either looked at from the point of view of the upstream neurons (efficacy) or the downstream neuron (contribution).

Wakefulness is associated with an increase of efficacy in both thalamocortical and intracortical synaptic contacts, which are significant but exhibit small to negligible effect sizes (Fig. 3b, avg. eff. TCA to BWN aw./ane. = 2.9 ± 0.91/2.5 ± 0.85 %, n = 68, p = 0.0014, δ = 0.27 (small); avg. eff. TCA to NWN aw./ane. = 3.29 ± 0.10/2.51 ± 0.07 %, n = 549, p = 2.9*10^-18, δ = 0.21 (small); avg. eff. BWN to BWN aw./ane. = 4.2 ± 0.47/4.05 ± 0.65 %, n = 152, p = 0.8, δ = 0.12 (negligible); avg. eff. BWN to NWN aw./ane. = 6.24 ± 0.2/4.86 ± 0.18 %, n = 552, p = 2.7*10^-17, δ = 0.21 (small)). In contrast, wakefulness induces significant but weak increase of contribution to BWN at thalamocortical synapses, meanwhile showing no change toward NWN, again with small to negligible effect sizes (Fig. 3b, avg. cont. TCA to BWN aw./ane. = 3.36 ± 0.33/2.68 ± 0.32 %, p = 0.00053, δ = 0.22, n = 68, TCA to NWN aw./ane. = 2.66 ± 0.069/2.85 ± 0.099 %, n = 549, p = 0.31, δ = 0.02 (negligible)). In parallel, wakefulness drives a general decrease of contribution in intracortical synaptic contacts, toward both downstream BWN and NWN (Fig. 3b, avg. cont. BWN to BWN aw./ane. =2.41 ± 0.49/2.59 ± 0.47 %, n = 152, p = 0.046, δ = 0.04; avg. cont. BWN to NWN aw./ane. = 1.75 ± 0.07/2.7 ± 0.12 %, n = 552; p = 1.4*10^-13, δ = 0.17; n = 11 mice). Altogether these results show that wakefulness is associated with relatively small changes of synaptic properties as they all have small to negligible effect sizes. It is interesting to notice that (1) wakefulness is associated with homogeneous increase of efficacy; (2) wakefulness has a differential effect on synaptic contribution, seemingly decreasing in intracortical synaptic contacts. This latter result further argues that intracortical downstream NWN are under the influence of other inputs during wakefulness. This pushes further the hypothesis that most NWN firings are led by TCA under wakeful conditions, meanwhile remaining less sensitive to their intracortical inputs despite their higher efficacy (Sup. Fig. 4a-b). To further examine these tendencies, we turned back to other quantifications of spikes transmission such as mutual information, here evaluated within each synaptic contact to evaluate the common entropy of the spike trains at both sides of the functional synaptic contact.

Classical correlation estimates such as Pearson correlation coefficient, or cross-correlation, can be ill suited when dealing with non-linear relationships between spike trains. To verify that we turned to mutual information, quantifying the common information based on common entropy between two spike trains (methods), actually used as an alternative to CCG detection when looking for synaptic contacts between neurons (Yamada et al., 1993, Jomaa et al., 2019). Given the strong correlation between firing rate and entropy in the spike train of single neurons (Sup. Fig. 7a-b), it is critical to normalize the obtained mutual information values to avoid biases from the change of firing themselves. In our case, we normalized mutual information by the entropy of the pre-synaptic spike train (London et al., 2002), then named proficiency which becomes unitless (Fig. 3 c-d). We observed in both awake and anesthetized states an unchanged and strong (apparently linear) relationship between efficacy and proficiency (Fig. 3c-d). The slight differences between downstream NWN versus BWN relationship remained similar in wakeful versus anesthetized conditions in both synaptic contacts (Wakeful TCA to BWN/NWN r = 0.99/0.84, anesthetized TCA to BWN/NWN r= 0.99/0.83, n = 68/549 synapses resp., n = 11 mice; Wakeful BWN to BWN/NWN r = 0.75/0.75, anesthetized BWN to BWN/NWN r = 0.80/0.74, n = 152/552 synapses resp., n = 11 mice). The apparent sharper relationship between efficacy and proficiency in anesthetized condition (Fig. 3d) is likely the reflection of the decreased efficacy during anesthesia (significant but no large effect size, see above, Fig. 3b). There is a stronger relationship between proficiency and efficacy in thalamocortical contacts (Fig. 3d) despite their lower synaptic efficacy (Fig. 3b, Sup. Fig. 5a-b), which suggests that proficiency is rather related to contribution (Fig. 3b, Sup. Fig. 5a-b). It nevertheless strengthens our claim that thalamocortical contacts, more than intracortical ones, are instrumental to the levels of activity of NWN.

Altogether, these results suggest that wakefulness is associated with small changes of synaptic properties and big changes of firing rates.

### Thalamocortical inputs set the tone of cortical inhibitory levels during wakefulness

To overlay these different results, we illustrate all wakefulness-associated changes in both thalamocortical (Fig. 4, left column) and intracortical (Fig. 4, right columns) contacts by reporting MLI of: upstream firing, downstream firing, efficacy, contribution, and proficiency (normalized mutual information).

First, when looking into meaningful changes induced by wakefulness, upstream firing and downstream firing show large effect size when compared to zeros in thalamocortical contacts (Fig. 4a, δ = 0.961/0.977, n = 617). By opposition, in intracortical contacts only the MLI of the downstream firing showed a large effect size (Fig. 4a, δ = 0.710, n = 704, n mice = 11). Second, when comparing MLI to each other, again upstream and downstream firing stand out in thalamocortical contact as the properties mostly impacted by wakefulness (δ of MLIs from resp. Upstr./Downstr. to Efficacies = 0.745/0.84; Upstr./Downstr. to Contribution = 0.867/0.916, Upstr./Downstr. to Proficiency = 0.753/0.828, n = 617). In addition, these two properties at thalamocortical synapses show neither medium nor big effect size between each other (δ MLIs Upstr. To Downstr. = –0.262, n = 617). Third, in intracortical contacts, there are barely any properties which show a large effect size when compared to zeros. Only the MLI of the firing of the downstream neurons exhibits a large effect size (δ = 0.710). Last, comparatively in intracortical contacts, most MLI compared to the downstream synaptic firing showed large effect size: MLIs of upstream firing, efficacy, contribution, and proficiency (δ to resp. Upstr./Eff./Cont./Prof. = –0.356/0.369/0.532/0.445). In either synaptic contact, comparing properties based on the downstream neuronal identity (either NWN versus BWN, red and black, Fig. 4a-b) showed no large effect size. The contribution of thalamocortical contact to NWN stands out as the most noticeable difference compared to their BWN counterparts (p = 7.3*10^-4, δ = –0.251, n = 549 NWN/68 BWN, n = 11 mice; other thalamocortical δ BWN vs NWN in resp. Upstr./Downstr. fir. /Efficacy/Pro. = –0.000/0.089/-0.048/0.073, or intracortical δ for BWN vs NWN in resp. Upstr./Downstr. fir./Efficacy/Contribution/Pro. = –0.012/0.203/0.11/-0.106/0.017).

Altogether these aligned results highlight how intracortical downstream firing stands out, compared to other intracortical synaptic properties, exhibiting similar wakeful modulations as observed at thalamocortical synapses. In addition, it is noticeable to observe how little synaptic properties are changed despite the strong changes of firing in all synaptic contact and conditions.

### Modelling connectivities only mimic biology with outnumbering thalamocortical inputs

To further assess the validity of our results, we used mathematical modelling to explore other connectivity schemes impossible to evaluate in *in vivo* settings. To do so, we adapted a delayed Hawkes Network (Hawkes 1971) to produce a simplified model of spike transmission in the visual pathway, composed of interconnected TCA, NWN and BWN in different numbers (Fig. 5). Our model simulates spike transmissions using the synaptic efficacies measured from our experimental data (methods, Fig. 5b), within a fixed window after receiving a presynaptic spike (0.8 to 2.8 ms, Fig. 5b). The TCA, BWN, and NWN populations are randomly connected with fixed probabilities (Fig. 2, Sup. Fig. 5). We simulated 9 thalamocortical connections per TCA, toward both BWN/NWN with a ratio of 10/90% (Fig. 2b), and 1.2 connections per BWN with a ratio of 20/80% toward resp. BWN/NWN (Fig. 2d). We also included inhibitory connections from NWN back to BWN (using a ratio of 10/90%, not imposing reciprocal connections, Sup. Fig. 5d). We used two efficacy values in thalamo– and intracortical connections as characterized in wakeful and anesthetized states (Fig. 3, Sup. Fig. 8a-b). We added a background firing rate in each neuronal population to account for: 1-the existence of other unmonitored inputs, and 2-the presence of spiking activity even in anesthetized conditions (Sup. Fig. 8c-d). Thereby, this modelling allows for the systematic probing of firing levels in TCA, BWN, and NWN populations, varying the numbers of thalamocortical and intracortical synaptic contacts, simply imposing a wakeful increase of activity from TCA only (Fig. 5a).

By first imposing the model to strictly follow our empirical connectivity observations, we tried using a ratio for the proportion of the different cell types of 10 TCA for 20 NWN and for 60 BWN, matching the captured populations (Fig. 1, n = 545 TCA/ 1272 NWN/ 3219 BWN, n = 11 mice). Using such a population ratio, we systematically failed to reproduce the wakeful increase of activity in NWN (Fig. 5d, (I)). Our (oversimplified) modelling is accounting for a very simple dynamic that NWN firing changes are reflections of the balance between their different synaptic inputs. In an oversimplified view, NWN MLI of firing is “pulled-up” by the wakeful increase of firing from TCA, and “push down” by the less changing BWN firings (Fig. 5d), smoothly integrating from the underlying synapses numbers from either source, as illustrated in the corresponding phase diagram (Fig. 5e). Consequently, we could only recover some wakeful increases of firing specifically in NWN when artificially increasing the numbers of TCA/NWN/BWN to 25/20/60 neurons (Fig. 5c-d, “Compensating connectivity”, (II)), thereby increasing TCA synaptic inputs to NWN from 80 to 225, meanwhile intracortical connections remain at 72 (Fig. 5a-b-c). This approximation was based on the fact that TCA amplitude is relatively small (∼ 40 µV peak-to-peak, Sibille et al., 2024, 73 ± 1.2 µV, n = 545, n = 11 mice, Sup. Fig. 3a), thereby likely under sampled in our recordings. Using this overestimation of 25 TCA instead of 10, we only could reach part of the observed MLI of firing (0.375 in the model vs 0.47 *in vivo*). To perfectly mimic our empirical observations, our model predicted that the numbers of TCA required would be 50TCA/20NWN/60BWN (Fig. 5d, “Extended TCA connectivity”, (III)), leading to a connectivity where NWN integrated on average from 9 thalamocortical versus 1 intracortical inputs. In addition, we also tested in our model a connectivity scheme without a preference toward NWN (Sup. Fig. 9a), which systematically failed to reproduce the MLI increase of firing of NWN, even when simulating 120 TCA to 60 BWN (i.e. 120*9 = 1080 thalamocortical connections, versus 60 BWN*1.2 = 72 connections, Sup. Fig. 9b-c).

Overall, this model quantifies the many possible connectivity schemes which strengthened the observation that the wakeful increase of firing of NWN being close to the TCA (0.47 vs 0.57, Fig. 1c-d) is possibly the reflection of an outnumbering number of upstream thalamocortical inputs of 9 to 1 versus upstream intracortical inputs.

## Discussion

### Summary

The presented results detail how wakefulness is associated with a specific increase of firing of both the populations of TCA and NWN, whereas comparatively the firing rate of BWN population minimally changes (Fig. 1). Such results can be accounted for by the specific connectivity from TCA to NWN and not BWN (90%/10%, Fig. 2). It should not be overlooked that thalamocortical downstream BWNs also exhibit a marked wakeful increase of firing (Fig. 3), unlike the rest of the BWN population (Fig. 1). These results suggest the presence of symmetric changes of firing at thalamocortical synapses regardless of the identity of the downstream neuron. The homogeneous increase of wakeful firing in NWN population further suggests that (1) our NWN sampling is homogeneous, likely from parvalbumin (PV) neurons given the localization of our tangential insertions, and (2) all captured NWN are under the influence of TCA inputs. These results thereby highlight the importance of feedforward modules at the first cortical relay of visual sensory processing, strengthening the importance of localized wakeful gain modulations (Ferguson et al., 2020), or center surround modulations (Keller et al., 2020), which are prominent during wakeful condition and disappear under anesthesia (Haider et al., 2013). In parallel, intracortical connectivity similarly shows a preference toward NWN (Fig. 2). Curiously, despite higher efficacies in intracortical connections (Fig. 3), wakeful increases of firing are only observable in intracortical downstream neurons and not upstream ones, with middle to large effect size (Fig. 4). This suggests an asymmetric modulation of firing at intracortical synapses, highlighting the predominant effect from other inputs, thereby arguing that TCA inputs (and not intracortical inputs) set the tone of wakeful NWN firing levels (Sup. Fig. 5). In addition, synaptic properties such as efficacy and contribution do not show middle to large effect sizes when comparing wakefulness to anesthetized conditions (Fig. 3-4).

### Modelling

Taken together, these results stress the key role of thalamocortical synaptic input in driving NWN activity levels in wakeful condition, unlike intracortical synaptic inputs, which is further supported by our mathematical modelling (Fig. 5). Indeed, through the different connectivity matrices we probed, our model indicated that only outnumbering TCA inputs with a ratio of 9:1 can mimic the observed wakeful NWN changes of firing. Given the previous reports of a larger physical spread of TCA (Sibille et al., 2024, Sup. Fig. 3b), and their larger diverging, converging connectivity scheme (Fig. 3), it could very well be that predominant TCA inputs are reflected in the true connectivity of the mouse brain. Indeed, previous experimental reports showed that up to 40 different thalamocortical inputs can be overlapping in the same cortical position (Jin et al., 2011), when others claimed up to 85 converging synaptic connections from thalamocortical inputs onto a single barreloid cortical neuron (Bruno et al., 2006). Alternative mechanism(s) could also account for the observed increases of firing, which are not included in our model, such as wakeful increase of excitability due to cholinergic inputs (Constantinople et al., 2011), changes of membrane properties, or dendro-somatic coupling (Suzuki et al., 2020), among the many other hypothesis underlying anesthetics effect characterized *in vitro* (rev. Platholi et al., 2022, Hao et al., 2020, Zhou et al 2012, Covarrubias et al., 2015). Patch clamp and morphological reconstruction have established that ground truth connectivity in the brain exhibits similarities and differences to our connectivity matrix (Campagnola et al., 2022, Potjans et al, 2014). These results are calling for a better understanding of non-linear spiking production and spike transmission, which can be further complexified by short-term plasticity (Teh et al., 2026).

### Normalized mutual information

Next, we looked into quantities such as normalized mutual information (Proficiency) which is thought to capture nonlinear regularities between spikes trains; to further confirm this differential influence from thalamo-versus intracortical inputs on wakeful cortical processing. Similar proficiency in both awake and anesthetized states strengthened the interpretation that wakeful firing levels are unrelated to changes of synaptic properties (Fig. 3-4). Comparing proficiency with efficacy strengthens the hypothesis that mutual information is a possible proxy for detecting synaptic contacts (Fig. 3, Yamada et al., 1993, Jomaa et al., 2019), which we complemented with quantifications in connected versus non-connected neurons (Sup. Fig. 7c-d). Nevertheless, the stronger relationship between efficacy and proficiency in thalamocortical contacts, compared to intracortical contacts (Fig. 3), further argues for different transmission properties in these two different contacts.

### GABAergic mechanism(s)

There is a long-standing breadth of knowledge in the field of anesthesia which already detailed the specificity of anesthetic actions on synaptic transmission (rev. Platholi et al., 2022, rev. Hao et al., 2020) and on ion channels (rev. Zhou et al 2012, Covarrubias et al., 2015). These results thereby feed several hypotheses on the mechanisms underlying the loss of consciousness. It is important to point out the classical assumption is that anesthesia is associated with a loss of activity, by way of over-activation of GABA transmission, which is here somehow paradoxical given the generalized reduction of NWN activity under isoflurane (Fig. 1). The possibility that isoflurane specifically enhanced GABAergic transmission in the thalamus (Muheyati et al., 2024) still remains unresolved. Direct thalamic pharmacological manipulation *in vivo* would potentially permit the reproduction of anesthetized effects without other possible co-modulations.

### Inferring mechanism(s) in vivo

Overall, it is important to discuss the specificity of anesthetic in the eyes of *in vivo* results, where the effect on the entire organ can be interpreted as unravelling the most sensitive location where the anesthetic acts (Leung et al., 2014), or even the origin of the observed effect (Cacciatore et al., 2025). For example, there is a discrepancy of effects in different brain regions when applying isoflurane, depending on whether one looks into the primary neocortex or elsewhere. For example, isoflurane induces a decrease of visual responses in V1 but induces an increase of auditory responses in A1 (Raz et al., 2014), when curiously isoflurane reduces A1 consequent bursting activity (Hentschke et al., 2017). Such longer effects can be associated with reports of a marked change of sensory responses organization under isoflurane in A1 (Filipchuk et al., 2022). More recently, it has been observed in S1 that anesthesia is associated with a general decrease of activity in Somatostatin-expressing (SST), PV and excitatory neurons (Sugino et al., 2026). On the other hand, in non-primary cortical areas, thalamocortical inputs seem barely impacted by isoflurane when corticocortical inputs are strongly changed (Murphy et al., 2019), thereby supporting the seminal idea of a loss of top-down modulations under anesthesia (Imas et al., 2005). This particular corticocortical modulation is probed in slices with electric stimulations in layer 1 (Murphy et al., 2019), thereby not overlapping with our experimental conditions where the intracortical connectivity we monitored is highly localized, as it rarely extends over more than 250 μm (Sup. Fig. 5). These separate results taken together ask that isoflurane effects on thalamocortical and corticocortical connections should be considered in a region-specific manner, and likely in a layer-dependent manner. Last, the thalamus is also under the influence of lower brain regions which can be strongly influenced by anesthetic, as shown in the auditory system, where the cochlear nucleus loses its tonicity even under a low dose of isoflurane (Gosselin et al., 2024).

When opposing the specific effect of anesthetic to the physiology observed during wakefulness, it is important to keep in mind that there are multiple additional unchecked confounding factors, such as neuromodulators in S1 (Constantinople et al., 2011), modulation from pure darkness in V1 (Pakan et al., 2016), or running (Niell et al., 2010), and facial movements (Stringer et al., 2019). In particular, body movements are associated with strong co-modulations of visual cortex activity, which are negligible in our experimental settings because they spontaneously occur during only 10 to 15% of our awake recording periods (Kala et al., 2026). It still remains to be explored how firing rates and synaptic properties are differently impacted during active behavior periods, versus visual ones.

### Inferring the mechanism(s) in our study

Still, there are several lines of question that remain open, in particular in regard to the mechanistic origin of the wakeful modulations we observed. Our results could be simplified as wakefulness is solely acting on TCA neurons, as implemented in our modelling (Fig 5). This mechanism somehow does not align with wakefulness modulation through the noradrenergic pathway (Constantinople et al., 2011), nor with the evidence that layer 5 pyramidal neurons are specifically impacted by different anesthetics (Bharioke et al., 2022), possibly in their dendro-somatic coupling (Suzuki et al., 2020). In addition, a few papers even report healthy wakeful modulations without thalamic inputs in sensory areas (Constantinople et al., 2011), meanwhile thalamic sectioning does not forbid sleep slow oscillations (Fuller et al., 2011). On the other hand, central and ventral thalamus are implied in wakeful processes, either in the recovery from anesthesia, or from sleep, highlighting the central role of the centromedian-parafascicular complex (Cacciatore et al., 2025). This particular brain region has quite some overlap with the paraventricular regions (Ren et al., 2018, Wang et al., 2023) underlying the ascending arousal system, which is still at play in recovering from sleep through loud sounds (Boccalaro et al., 2025). We do not intend to claim that the change of TCA activity in the visual system is the leading factor driving wakefulness, but rather that these changes of activity are associated with wakeful and anesthetized states. To do so, one could hypothesize larger interacting loops, such as: a loss of TCA activity => loss of NWN activity => loss of excitation/inhibition balance => gain of synchronicity within the cortex (Bharioke et al., 2022), and the loss of dendro-somatic coupling (Suzuki et al., 2020). Following such a hypothesis would then suggests an anesthetic origin in either thalamic inhibitory neurons (Muheyati et al., 2024), or layer 6 CT neurons (Hönigsperger et al., 2024), not accounting for specific wakeful norepinephrine and acetylcholine activations (Constantinople et al., 2011).

## Conclusion

One cornerstone in this study lies in this newer capability offered by tangential Neuropixels probe insertions to harness parallel synaptic contacts simultaneously (Fig. 2 & Sup. Fig. 5-6); here focused on thalamocortical and intracortical synapses (Fig. 2). This particularity sets apart connected neurons from the surrounding populations. This is exemplified by the strong discrepancy we observe between the population of BWN which apparently does not change their firing during wakefulness and the sub-populations of thalamocortical downstream BWN which exhibit a strong wakeful increase of firing. These altogether argue that first, TCA set the tone of NWN wakeful firing, second outnumbering TCA connections to NWN may underline the homogeneous increase of wakeful firing in NWN, and last intracortical connections are less impacted by wakefulness; altogether suggesting different wakeful tonicity in thalamocortical and intracortical networks.

## MATERIAL AND METHODS

### Animals, surgery, insertion, and staining

All data presented stem from head-fixed experiments (n = 11), which protocols were acknowledged beforehand by the local authority (Landesamt für Gesundheit und Soziales Berlin – G 0142/18). Maximum care was taken to decrease the number of used animals; all adult C57BL/6J male mice originated from local breeding facilities (Charité-Forschungseinrichtung für Experimentelle Medizin). A first surgery was performed to construct a head post, implanted ∼ 2 weeks prior to the recording day. This implantation was realized in a stereotactic frame (Narishige) under isoflurane (2.5% in oxygen Cp-Pharma G227L19A). Painkiller (metamizole 200 mg/kg, Zentiva-Novaminsulfon) was provided in the drinking water 1 day prior to, and 3 days post-implantation, renewed daily. During surgery, temperature was constantly monitored (FHC-DC), eye ointment (Vidisic) was applied to avoid drying of the eyes. The head-post was fixed to the skull by way of building a large dental cement-based crown (Paladur, Kulzer). This crown ranged from bregma to post-cerebellum skull surfaces, extending laterally on both lateral ridges to stabilize the implant. The crown walls were made of equal height to produce a well above the head of the animal, thereby providing a grounding liquid bath during recording. Too high walls should be avoided to allow for the flat tangential cortical insertion, as done previously for both tangential cortical (Sibille et al., 2024) and superior collicular insertions (Sibille et al., 2022b; Gehr et al., 2023). According to protocols, mice were gradually habituated to head fixations on the set-up, increasing exposure time from 30 s to 2 hours. During habituation, classical visual stimuli, i.e. sparse noise, moving bars, moving gratings and full-field chirp stimuli were exposed to reduce putative stress during recording day. A second surgery was done in the morning prior to data acquisition to produce the craniotomy, leaving a recovery period of at least 2 hours before recording. The craniotomy was performed with the following coordinates: 0 to 900 μm AP, +250 to –250 μm ML, and –100 to –500 μm DV from lambda, carefully avoiding the midline blood vessel. It is advised to leave a bit of skull above it, cautiously polishing the skull opposite to the insertion point in order to facilitate the flat tangential insertion. In the meantime, an extra grounding wire was added within the crown to reduce the noise levels. Extra care should be given to grounding when attempting to measure TCA, which have small amplitudes (Sup. Fig. 3a, Sibille et al., 2024). During recording, the crown was regularly refilled with PBS to keep electrical noise as low as possible.

For our acute recording, the animal was either placed on a fixed platform (n = 7) or a running band (n = 4) to proceed with flat medio-lateral tangential insertions in the visual cortex. These insertions are done by placing the probe and micromanipulator arm (New Scale) within a coronal plane at a –10 to –15° angle from the azimuthal/interaural plane. The arm should come from above to avoid obstructing the visual field of the animal. Then, according to space and crown height, the probe tip is placed onto the brain tissue surface, as close as feasible from the midline, to be then inserted for at least 4-5 mm through the cortical mantle (Sup. Fig. 2d). The insertion is ended by a withdrawal of 50-100 µm to release accumulated mechanical pressure. To avoid bending of the probe from the water tension surface, the PBS solution should be removed prior to probe placement. It is important to notice that preliminary recordings are done to perform 1-semi online detection of receptive field mapping (https://github.com/KremkowLab/tangential_recording, Sibille et al., 2022b) and 2-semi online TCA detection (https://github.com/KremkowLab/Cortical-Axon-on-Neuropixels-in-Kilosort, Sibille et al., 2024). It is important to obtain visually driven activity where TCA are detected. A good recording should ideally contain at least 50 TCA detected in the preliminary recording, which can be done on the receptive field mapping pre-recording (lasting ∼ 5 min). Furthermore, it can be difficult to get concomitant thalamocortical and intracortical connectivity within a single tangential insertion (Fig. 2, Sup. Fig. 6), due either to a placement more exclusively in layer 4 capturing TCA inputs (Sup. Fig. 6a), or slightly higher in layer 2-3 capturing intracortical connectivity (Sup. Fig. 6b). In case of doubt, such as poor visual activity, or poor TCA reach, the probe should be re-inserted, aiming for a better location. It is to be noticed that 25-30% of the animals being conditioned do not convey a good recording, which thereby reduces post-recording analysis. This rejection is mostly based on the preliminary recording evaluation, where visual responses are not present, or TCA are too few.

Once a satisfying location is obtained, the probe is left to settle in the tissue for ∼20-30 minutes before recording can be started and visual stimuli exposed. After the first list of visual stimuli were exposed (∼1 to 1.5 h), the anesthetic mask (1% Isoflurane) was then slowly approached to anesthetize the mice while keeping the recording running, allowing for later paired comparison between wakeful and anesthetized states (n = 11 mice). Due to unknown reasons, we had a mouse sustaining anesthesia in a pseudo wakeful state, characterized by facial and body movements throughout the anesthesia period, leading to rejection from further analysis. Once the recording was finished, the Neuropixels probe was partially removed (leaving the tip in the tissue), coated with DiI (Abcam-ab145311, in EtOH), and reinserted in the same location. If DiI coating is done before the first insertion, repositioning of the probe after preliminary recordings will impair later reconstruction. Mice were terminated with an excess of Isoflurane (>4%), allowing for subsequent cardiac perfusion, first with PBS, then with paraformaldehyde (PFA, 4% in PBS). Once the brains are extracted, they are left in PFA overnight to be sliced (100 µm) and mounted (DAPI-Fluoromount-G Biozol Cat. 0100-20) within the next day(s). Mounted slices were pictured under microscope to allow for 3D reconstructions using SHARP track (Shamash et al., 2018) for each channel to be relocated with the Allen Mouse Brain Common Coordinate Framework (Sup. Fig. 2).

### Software, hardware, and visual stimuli

Neuropixels probes (1.0) signal was measured with Open-Ephys acquisitions software (www.open-ephys.org) plugged in a PXIe system (National Instrument NI-PXIe-1071). Standard cluster quality metrics were calculated to remove units of bad quality (cf. below), where the manual curing was performed based on Kilosort 2.5 output. All data analysis was performed in Python 3 (www.anaconda.com).

Statistical testing was done using two-sided Wilcoxon rank-sum (for unpaired comparison between two distributions), or Wilcoxon signed-rank test (when doing paired comparison, Fig. 1e, 2f, 3b-d), or one-sample Wilcoxon signed-rank test (when evaluating the changes through the MLI, comparing to a distribution of zeros of similar size); p-values below 0.001/0.01/0.05 are indicated by ***/**/* respectively. Modulation indexes were evaluated for the firing rate, efficacy, contribution, and proficiency 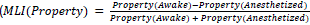, Fig. 1-4). All script and analysis were done in home-made script (numpy, scipy, pandas, sklearn), except for the spike sorting done under Matlab. Standard errors of the mean (SEM) are being used throughout the manuscript when reporting means quantifications, unless otherwise stated. The strength of the effect of comparison was further evaluated with the Cliff delta value (<0.11: negligible; 0.11-0.28: small; 0.28-0.43: medium; 0.43<: large). Medium to large Cliff delta size effects were only reported in the figure with a “δ” sign. Given two distributions of respectively size *m* and *n* with the values *x_i_* and *x_j_* then *Cliff δ* = (∑*_i_*_,*j*_ [*x_i_* > *x_j_*] − [*x_i_* < *x_j_*])/*mn*. Visual stimuli were exposed using a monitor screen (Dell, calibrated at refresh rate = 120 Hz, mean luminance = 120 cd/m²). They cover alternating checker mapping (10°, 1 s), two sparse noises (dark or light targets, 15° squares, 2 targets/frames which last 100 ms, 20 repeats for each position on a 36 x 22 grid, cf. https://github.com/KremkowLab/tangential_recording), moving bars (white bars on black background, 10° in width, 12 directions with a fixed speed of 90°/s), and full-field chirp stimuli. This latter stimulus was produced as such: 1 s gray background, 2.2 s black, 3.3 s white, 3.3 s black, 2.2 s gray, followed by 8.75 s of sinusoidal intensity modulations at full contrast having increasing frequency (0.5 Hz to 11 Hz), followed by 8.75 s of a gradual increasing contrast (0 to 100%) at 0.4 Hz, which was ended by 2.2 s gray background; the entire sequence was repeated 10 to 20 times. For time alignment purposes, each visual stimulus frame (or start for the chirp) was accompanied by a transistor-transistor logic (TTL) pulse sent from the stimulus computer into the PXI system that is connected to the NI-DAQ (PXI-8381) of the Neuropixels recording PXIe for later time alignment.

### Spike sorting

Importantly, TCAs have been shown to be of thalamic origin as detailed in our previous paper with due controls and independent analysis (Sibille et al., 2024). First, TCAs are resistant to local application of muscimol. Second, TCA activity is abolished upon TTX injection in the dLGN. Third, TCA waveforms can be extracted in multiple Neuropixels insertions from the Allen openscope dataset, when comparatively corticocortical axonal arbor waveforms are not observable. Fourth, upon electric stimulation applied in the dLGN, TCA are firing before V1 neurons (Sibille et al., 2024). Last, when illustrating the different ISI exhibited by TCA, clear bursting patterns appear under anesthesia (Sup. Fig. 3c), similarly to previous observation in the thalamus of anesthetized cats (Reinagel et al., 1999). In this current paper, TCA are detected from Kilosort 2.5 (Fig. 1, Sup. Fig. 2), which we think is possibly ubiquitous to most other cortical areas and mammalian species which receive thalamocortical axons. (rev. Sibille et al., 2026). Vertical insertions also have the capability to capture TCA. We recommended using a dedicated change of KS 2.5, as made available online (https://github.com/KremkowLab/Cortical-Axon-on-Neuropixels-in-Kilosort).

Manual curation was done post-hoc using Phy2 (https://github.com/cortex-lab/phy/). It is important that the clusters with multiple temporal peaks and large spatial spreads are not removed during the manual curation step(s) because these features are two key properties distinguishing TCA (Sup. Fig. 3a, Sibille et al., 2024) from other normal V1 neuronal waveforms (rev. Sibille et al., 2026). Furthermore, often due to the smaller amplitudes of TCA waveforms (Sup. Fig. 3a, TCA amplitude = 73 ± 1.2 µV) certain quality controls such as amplitude cutoff, or even the MUA label given by KS, should be ignored first during preprocessing steps in order to be reexamined later. Once manual curation on Phy is finished, a second step of selection is done from recalculated waveforms and quality metrics. First a manual removal of double-counted spikes is done before scrutinizing inter-spike-interval violations (Sibille et al., 2022, > 0.05%). Isolation distance (> 10 a.u) was also used to remove bad units. Then recalculated waveforms are used to screen single-unit properties individually within a custom-written Graphical User Interface. This step is done to distinguish and label neuronal somatic waveforms, TCA waveforms, from putative unknown waveforms (Sibille et al., 2026). The recalculated waveforms were produced by averaging the raw Neuropixels signal for all spike times for each individual single-unit (up to 50 000 spikes). These recalculated waveforms also include compensations for the slight offset correction of 2.78 μs between neighboring channels within the same analog-digital-converter (Sibille et al., 2022). Last, an additional step was performed to compare the waveform from each cluster in wakeful versus anesthetized conditions. If any waveform discrepancy was observed between wakeful and anesthetized conditions, then the corresponding single-units were completely removed from the dataset and any further analysis. To distinguish between broad waveform (putative excitatory) neurons versus narrow-waveforms (putative inhibitory) neurons a threshold was fixed at 0.43 ms of their peak-to-trough duration on the waveforms at its “best channels” (with the biggest amplitude, Sup. Fig. 1), somehow likely sub-selecting PV and SST neurons (Valero et al., 2025).

### Receptive fields, firing properties and mutual information

For RFs measures, first the peri-stimulus-time-histogram (PSTH) was obtained by summing all activity on each exposed pixel over the entire stimulation duration (sparse noise). RF contours were interpolated using the 2D-cubic-interpolation function from SciPy packages, keeping there only clusters having a big enough SNR (1/std (RF) > 15, Sup. Fig. 2 f).

The neurons having inter-spike intervals of less than 5 ms were considered bursty. The burstiness was computed as the ratio of bursty spikes to all spikes (*n_spikes_*(*ISI* < *5 ms*)/*n_spikes_*(*all*)), Sup. Fig. 4 c-d). To estimate neuronal firing correlations, first the firing of every neuron was normalized between 0 and 1. Then an average firing of all cells was calculated to be used to evaluate its correlation to each individual neuronal normalized firing (Bharioke et al., 2022, Sup. Fig. 3 a-b). The coefficient of variation of firing of a cell is calculated as the standard deviation of its firing over time, divided by its average firing rate (Sup. Fig. 4 c-e), which quantifies the fluctuation of the neuronal activity over time (Bharioke et al., 2022). In addition, using Poincaré plots (ISI(n) vs ISI(n+1)) can reveal highly organized bursting firing patterns (Sup. Fig. 3c) observed only in a subset of TCA, and being absent in NWN or BWN.

Mutual information (bit) was estimated using homemade script (Fig. 3-4, Sup. Fig. 7). The entropies of either the pre-synaptic neurons (H0), the post-synaptic neurons (H1), but also the common entropy between spikes trains (H(0,1)) were used to calculate on both sides of a given detected functional connection the shared entropy (Fig. 3c-d, Fig. 4). The entropy of a spike train was approximated as the cumulative sum of the probability of each possible spiking state divided by the total number of possible states, as usually done when calculating the information content of a discrete time series. A window of 3 ms we use to approximate the different presence of a spike (1) or its absence (0), within the 3 ms window (Fig. 3, London et al. 2002), for all windows position (i).

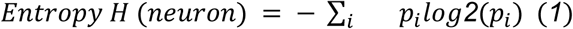

Consequently, the common entropy between two spikes trains are stemming from similar equations quantifying as either 1 when both spikes trains are in the same state (either both non firing or both firing in the 3 ms windows), or when they are in a different state.

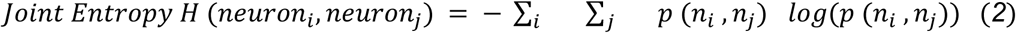

To estimate proficiency the obtained mutual information was divided by the pre-synaptic entropy thereby avoiding any effect for the changes of pre-synaptic firing (London et al. 2002). To estimate proficiency in locally unconnected pairs, only pairs of neurons within a 200 µm distance were chosen randomly from the unpaired neighboring neurons of the different types (Sup. Fig. 7b). Proficiency was estimated on the spikes train in either wakeful or anesthetized conditions.

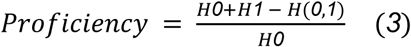

### Functional synaptic connectivity, detection and quantification

Connectivity in V1 between TCA, NWN, and BWN was detected using cross-correlation analysis (CCG, using pycorrelate https://github.com/tritemio/pycorrelate). Only neurons that have more than 2000 spikes been included for connectivity analysis. For each pair of neurons, a spike jitter was calculated in order to remove putative common correlations throughout the entire recording, which was done by randomizing every spike time within a 10 ms window. This jittered CCG was then subtracted from the raw CCG for monosynaptic detection (Fig. 2-3-4, Sup. Fig. 5). Four different types of connections were identified on the basis of significant peaks within different time windows: 0.5 to 3 ms positive peaks for monosynaptic excitatory connections, (with a minimum of 4 consecutive time bins), either from TCA to V1N for thalamocortical connectivity, or from V1N to V1N for intracortical connections (Sup. Fig. 5). In addition, central peak +-0.7ms was carefully scrutinized to isolate a symmetric peak for common inputs versus asymmetric ones followed by a large inhibitory trough for putative inhibitory connections (Sup. Fig. 5), as previously done in other cortical areas (Fujisawa et al., 2008). We identified inhibitory connections only when the CCG contained a combination of a central peak together with a significant decrease of correlated activity below the jitter correction by combining positive peak detection to negative peak detection in a flipped CCG within the +1 to 5 ms windows. For inhibitory connections, we imposed at least 8 consecutive time bins to remove possible noisy correlations within pairs of neurons. A peak was considered meaningful if its size reached at least 3 x the standard deviation of the CCG – the jittered CCG, as previously done (Sibille et al., 2022, Gehr et al., 2023). These settings for the detection of inhibitory connections may very well overlay with reciprocal connections, which is not the scope of the current paper solely based on excitatory functional connected pairs. Importantly, the detection of a connected pair was done using all spikes in both wakeful and anesthetized conditions. Once a connected pair was detected, different properties were extracted such as the relative distance (by the Euclidean distance between their respective best channels), their peak time (Sup. Fig. 5). Diverging and converging monosynaptic contacts were quantified as counting connections from one to several neurons and vice versa.

To estimate synaptic efficacy and contribution, the baseline of the CCG was used and subtracted from the CCG, instead of the jitter correction (Fig. 3a, Gehr et al., 2023), to avoid noisy estimations of synaptic strength. Efficacy was defined as the area under the monosynaptic peak of the baseline corrected CCG, divided by the total number of synaptic upstream spikes. To estimate the contribution, the reversed (post-pre) CCG was used instead, subtracting its baseline to quantify the peak area divided by the downstream neuronal firing (Fig. 3a). To compare synaptic properties (such as efficacy and contribution) in anesthetized versus wakeful conditions, CCGs had to be re-calculated from the spikes in each condition, to re-evaluate synaptic strengths using the very same method. It is important to notice that detecting synaptic contact by way of CCG analysis separately in awake versus anesthetized periods is unadvised because during anesthesia, firing rates can drop so strongly that barely any single-unit will pass the criterion of having more than 2000 spikes; leading to a strong under-estimation of the thalamocortical connectivity for example.

### Model setup

The network model is a variant of delayed Hawkes network (Hawkes 1971). It consists of 0 to 110 excitatory TCA inputs (the baseline should be chosen at 10, Fig. 5 is plotted with 25 TCA), 0 to 210 excitatory cortical BWNs (the baseline being chosen at 60 BWN), and 20 inhibitory cortical NWNs. These ratios have been chosen to match the population in our dataset (545/1272/3219 ⇔ 10/20/60). The neurons are modeled as Poisson neurons, with their spike distributions following a Poisson distribution with predefined baseline firing rate *λ_baseline_*. All neurons have refractory period of 1 ms and synaptic delay *τ_delay_* = *0*.*8* ms. For connected neuronal pairs, the postsynaptic neuron will have an elevated firing rate *λ_elevated_*(*t*), in addition to its baseline firing rate, after receiving a presynaptic input spike. More specifically, the elevated firing rate is the sum of firing rates induced by all presynaptic neurons at time *t*:

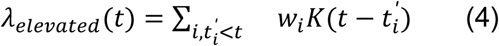

where *w_i_* is the *i*-th presynaptic input weight (efficacy), *t^′^* represents spike times of the *i*-th presynaptic neuron and

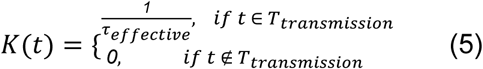

where *τ_effective_* = *2* ms is the duration of spike transmission and *T_transmission_* = [*τ_delay_*, *τ_delay_* + *τ_effective_*] ms is the effective spike transmission window. The spiking probability at time *t* is given by

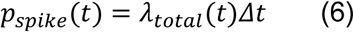

Where *λ_total_*(*t*) = *λ_baseline_*(*t*) + *λ_elevated_*(*t*) is the total firing rate and *Δt* = *0*.*1* ms is the temporal resolution of the simulation.

#### Model connectivity

The excitatory connections between different neuron populations (and the inhibitory connections) were all approximated from our empirical data (Sup. Fig. 5a-b). In the three types of connections: thalamocortical, intracortical, and inhibitory connections, all connection probabilities were fixed to the number of presynaptic neurons which were simulated. Specifically, we imposed on thalamocortical connections: 1– a fixed ratio of post-synaptic neurons of 90% toward NWN and 10% toward BWN, 2–9 post-synaptic connections per TCA (Fig. 2b), 3– a fixed efficacy of 2.9/2.5% to BWN and 3.29/2.51% to NWN in resp. wakeful and anesthetized conditions (Fig. 2f). In intracortical connections, we fixed: 1– the ratio of connections to 80% of NWN and 20% of BWN, 2-1.2 connections per BWN, 3– a fixed efficacy of 4.2/4.05% to BWN and 6.24/4.86% to NWN in resp. wakeful and anesthetized conditions (Fig. 2f). In inhibitory connections: 1– a fixed ratio of post-synaptic neurons of 90% toward NWN and 10% toward BWN, 2-2 post-synaptic connections per NWN (Sup. Fig. 5d), 3– a fixed efficacy of 5% to both BWN and NWN in wakeful and anesthetized conditions. Given such settings, varying TCA and BWN neuron numbers permits to automatically scale the model size up and down, co-varying the number of synaptic contacts, to simulate context impossible to test experimentally. In doing so we kept the imbalance between connection to NWN versus to BWN in the first set of simulations (Fig. 5, Sup. Fig. 8). Then we probed other possible connectivities, such as unspecific connectivity matrix (Sup. Fig. 9), where the ratio of downstream BWN/NWN are similar in all thalamocortical, intracortical and inhibitory contacts. These simulations completely failed to account for the wakeful increase of firing in NWN, despite increasing TCA numbers drastically (Sup. Fig. 9b-c).

### Code availability

Online analysis to calculate the visually driven MUA activity, equally as TCA pre-observation have been published in separate publications (Sibille et al., 2022, Sibille et al., 2024). Corresponding scripts are all available online on the lab GitHub pages and will be gladly shared to any interested scientists. Data sharing can also be done upon reasonable requests to the corresponding author.

### Funding

This study was funded by the German Research Foundation Deutsche Forschungsgemeinschaft (DFG) KR 4062/4–1, KR 4062/4–2, KR 4062/5-1 and KR 4062/6-1 (JK).

## Acknowledgements

The mouse head schema used in Fig. 1 a, and Sup. Fig. 2 a was adapted from https://doi.org/10.5281/zenodo.3925903. We thank Yota Kawashima for the help in analyzing mutual information. The manuscript was written with inputs from Francois David, Gregory Knoll, Yota Kawashima, and Bernardo Andrade.

## Author Contributions

Conception and design: J.S. and J.K.

Acquisition of data: J.S.

Analysis and interpretation of data: J.S., K.T., J.A.P.

Development and implementation of model: K.T., J.A.P.

Drafting or revising the article: J.S., K.T., J.A.P., J.K.

## SUPPLEMENTARY MATERIAL

**Supplementary Figure 1:**
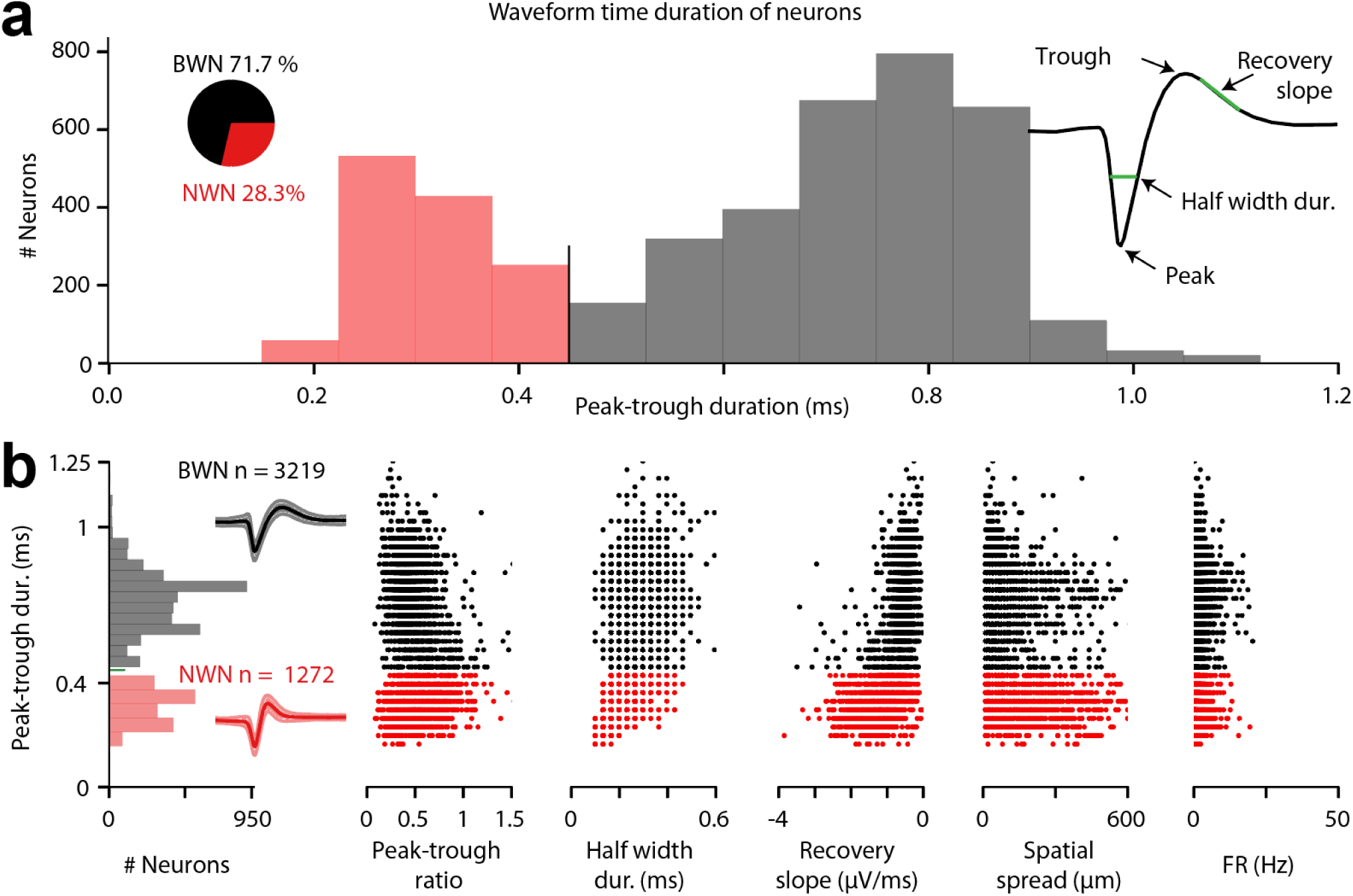
Broad and narrow-waveform neurons are distinguished by their peak-to-trough durations. **a**, Histogram of the different peak-to-trough values in all cortical neurons from the different tangential insertions (n = 11 mice), distinguishing 3219 BWN versus 1272 NWN. **b**, Classical properties from single channel waveforms aligned to the peak-to-trough histogram, in the corresponding order: peak-to-trough ratio, half width duration, recovery slopes, spatial spread, and firing rate. Averaged normalized waveforms are illustrated in the plot on the left, SEM between normalized waveforms is represented by the shaded area.

**Supplementary Figure 2:**
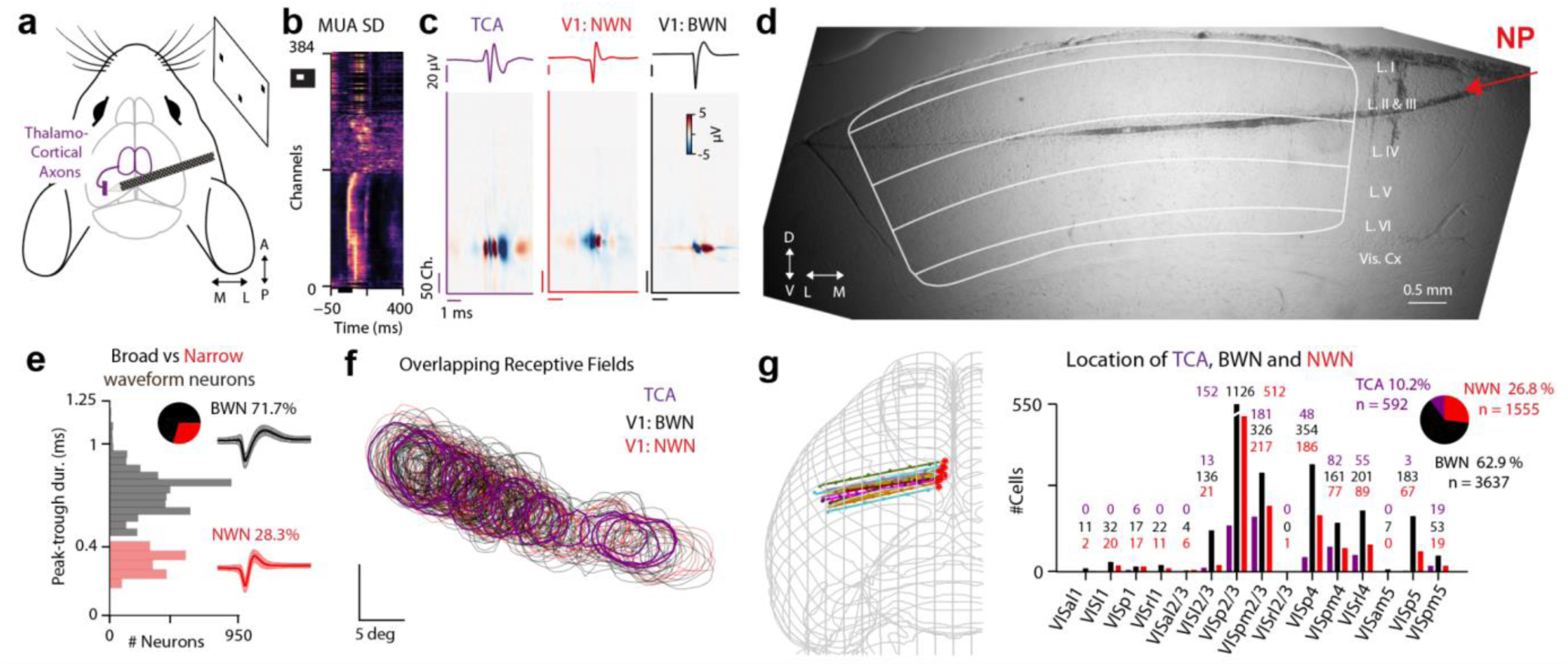
Layer locations and receptive fields from TCA, NWN, and BWN. **a**, Schematic of the tangential insertion. **b**, Visual responses obtained from sparse noise white square on black background inducing multi-unit-activity (MUA) over different channels of the probe. **c**, Typical multiple-channel-waveforms of the different thalamic axons (TCA, left, purple), NWN (middle, red) and BWN (right, black). **d**, Histological picture of a tangential insertion from the top, going through layer 2-3, layer 4. **e**, Histogram of the different duration of the peak-to-trough of the different NWN and BWN. **f**, Overlapping receptive fields stemming from TCA, NWN, and BWN captured in similar tissue location, spanning over different cortical retinotopic locations. **g**, Sharp-track reconstruction of the location of the different probe’s insertions in the different cortical layers to identify the layer’s location of the neurons throughout the dataset.

**Supplementary Figure 3:**
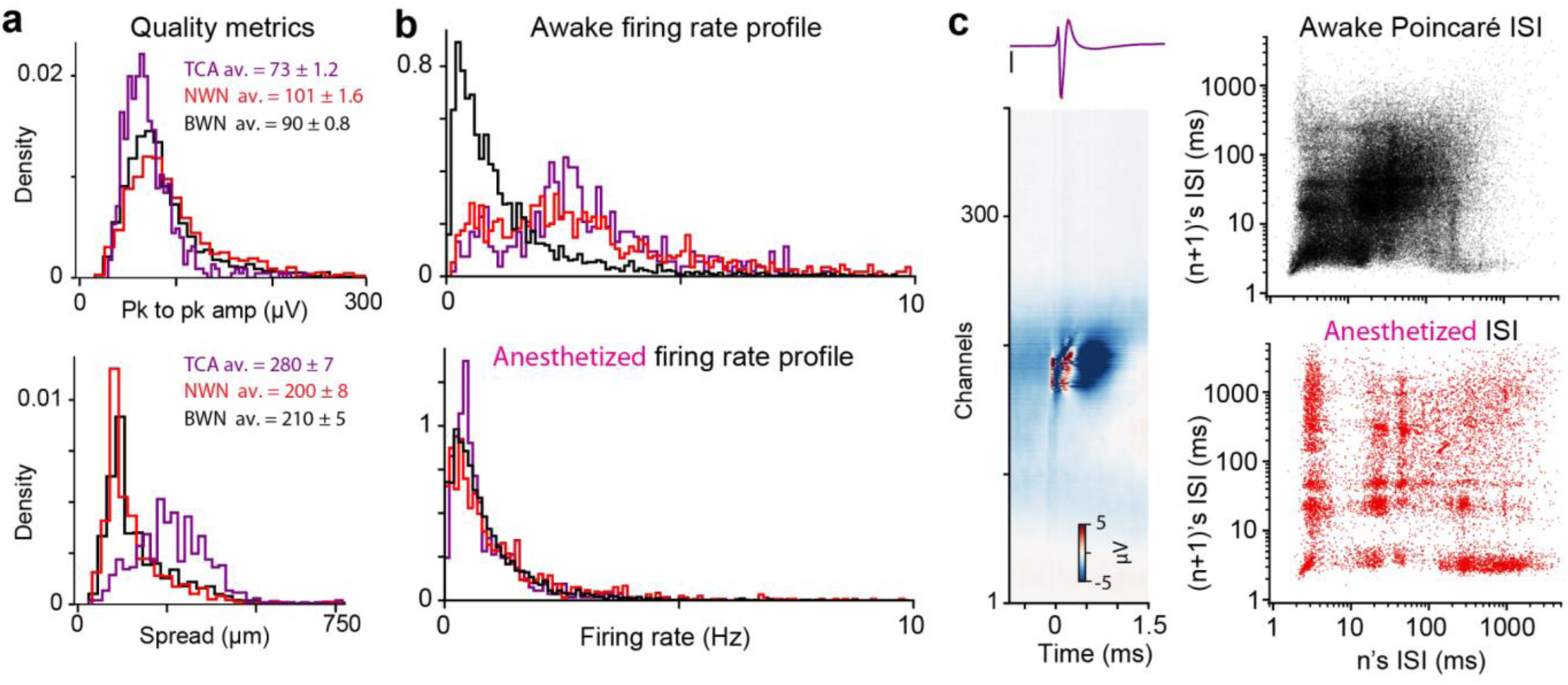
Firing properties of TCA, NWN, and BWN in awake versus anesthetized states. **a**. Quality metrics: Histogram of the single-unit amplitude (top, quantified in their peak-to-peak amplitude) and spread (right, quantified from the recalculated multichannel waveforms in micrometers). **b**. Histogram of the firing rate during awake (top) and anesthetized periods (bottom). Note that the apparent specific loss of high firing rate is mostly impacting TCA and NWN. **c**, Illustration of a TCA multichannel waveform (left, with its single channel profile on top, scale bar: 20 µV), and the Poincaré plots of the spikes inter-spikes-interval (ISI) in awake (top) and anesthetized (bottom) conditions. Note the structures appearing in anesthetized conditions suggesting more bursty firing patterns. These structures are observed on rare occasions, and apparently not present in NWN nor BW.

**Supplementary Figure 4:**
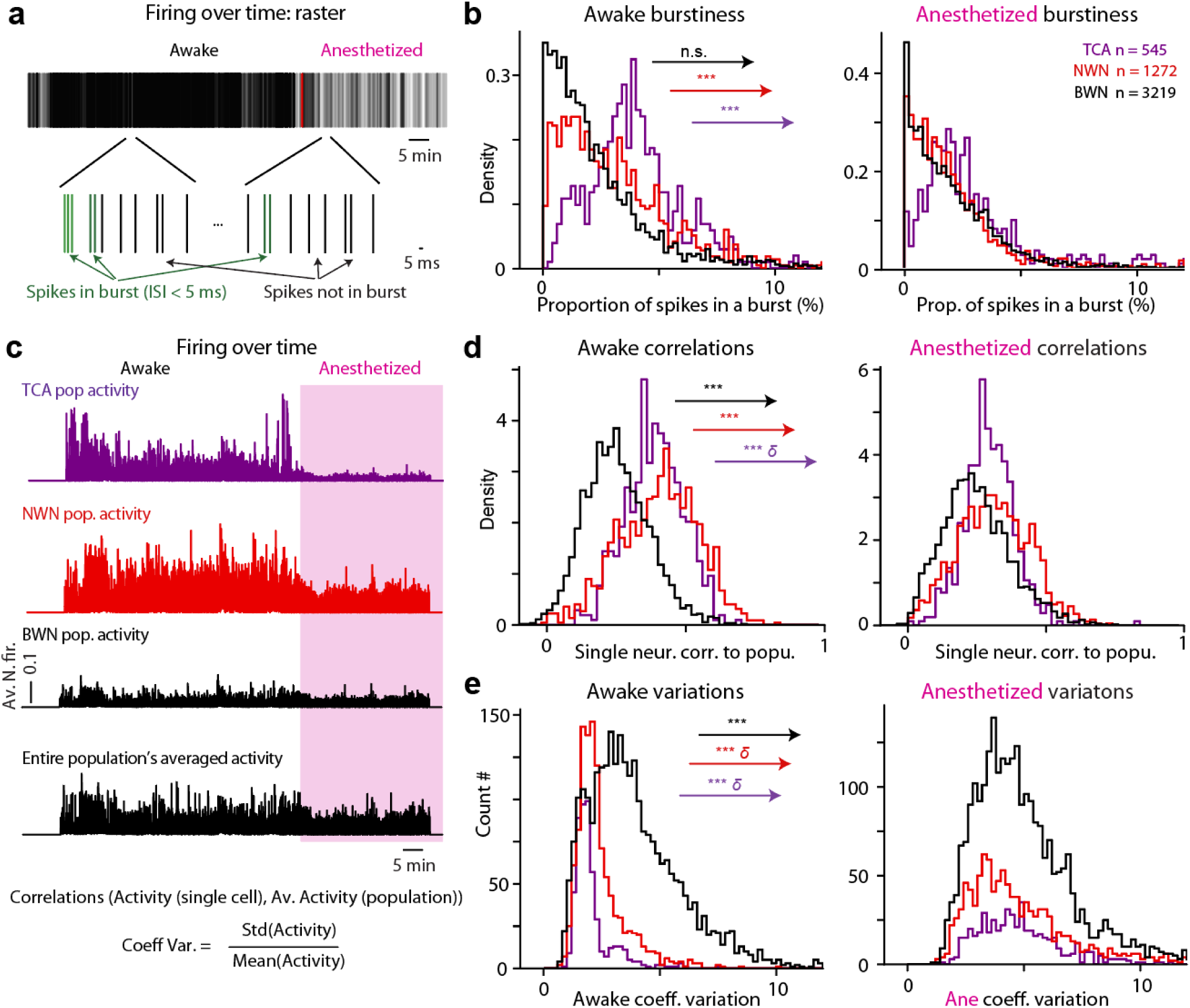
Firing properties of TCA, NWN, and BWN in awake versus anesthetized states. **a**, Schematic illustration of the timing of firing of a given neuron (top) to illustrate the quantification bursty versus non-bursty spikes, based on inter-spike-intervals above or below 5 ms. **b**, Histogram of the percentage of bursty spikes in awake (left) and anesthetized (right) periods (Aw. vs ane.: TCA p = 1.12*10^-11, δ = 0.25, NWN p = 3.43*10^-88, δ = 0.25, BWN p = 0.25, δ = 0.013). **c**, Schematic of the profiles of activity used to characterize both the coefficient of variation and the correlation to the population. The entire population averaged activity (bottom row) is to be compared to each single neuron activity profile individually, to quantify its correlation to the population. **d**, Histogram of the correlation values of TCA, NWN, and BWN to the population activity during awake (left) and anesthetized (right) periods (Aw. vs ane.: TCA p = 7.8*10^-69, δ = 0.5, NWN p = 3.17*10^-111, δ = 0.32, BWN p = 3.24*10^-6, δ = 0.02). **e**, Histogram of the coefficient of variation of TCA, NWN, and BWN during awake (left) and anesthetized (right) periods, estimated from normalized firing rate profiles (Aw. vs ane.: TCA p = 2.5*10^-90, δ = 0.88, NWN p = 1.7*10^-194, δ = 0.7, BWN p = 1.62*10^-138, δ = 0.27).

**Supplementary Figure 5:**
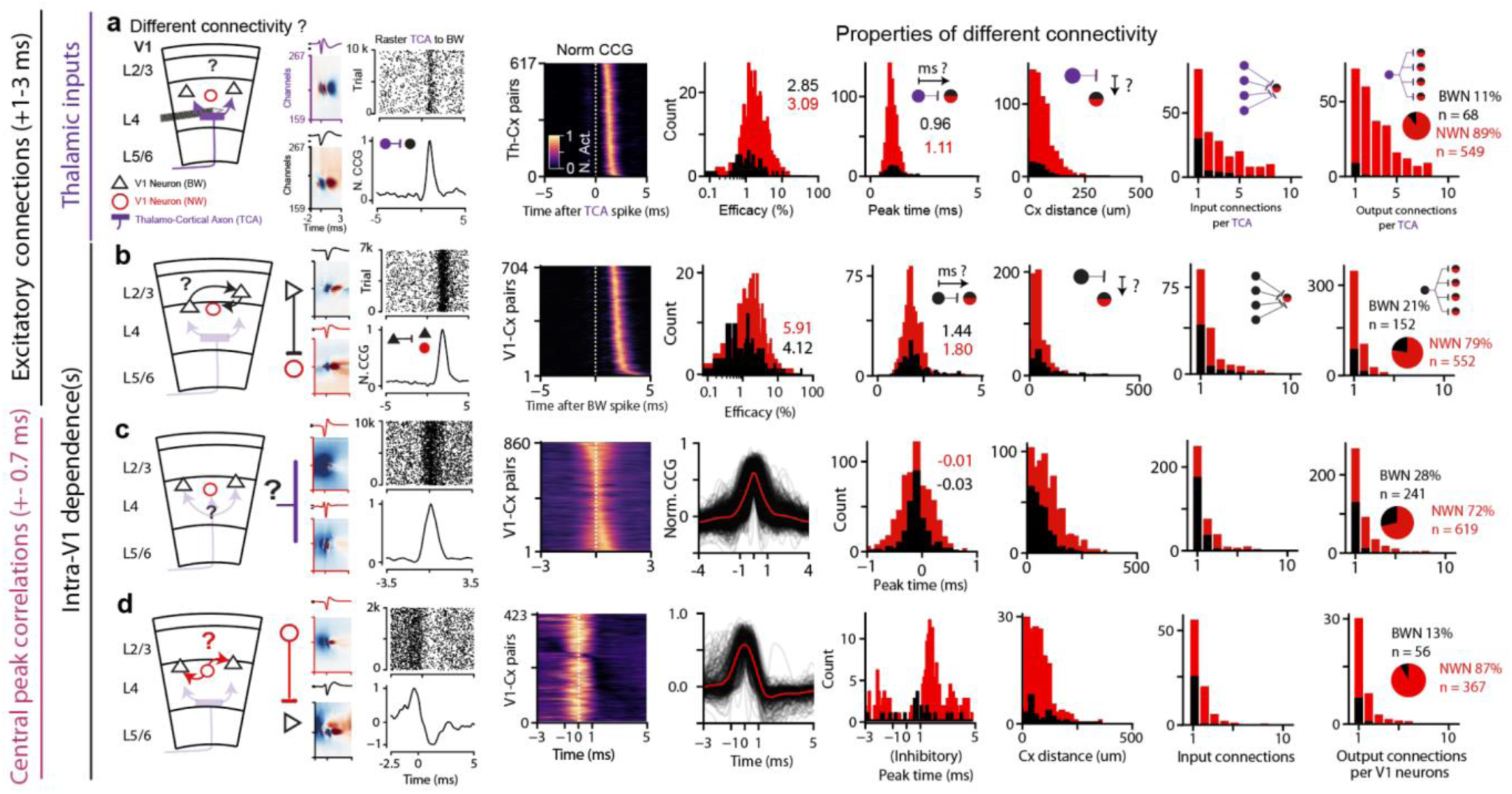
Overview of the 4 different types of CCG pattern extracted in tangential recordings. **a-b**, Repeat of the details about the monosynaptic excitatory functional synaptic contacts that are analyzed throughout the paper (n = 11 mice). **c-d**, Two other types of putative mono-synaptic interactions detected with a prominent central peak. **a**, Thalamic excitatory monosynaptic inputs, schematic of the connection type between TCA and either NWN or BWN (left). Details of the different properties of thalamic monosynaptic pairs (left to right): overlapping normalized CCG, efficacy, peak time, physical distance between upstream– and downstream synaptic pairs, converging, and diverging properties. Note that these thalamic inputs provide large connections over distances above 150 µm and can connect to up to 8-9 different post-synaptic neurons. Furthermore, thalamic inputs connect mostly to NWN (89%). **b**, Similar plotting for intracortical excitatory monosynaptic pairs. Note that again most of these connections are toward NWN (79%). **c**, Details of the different properties of common inputs: overlapping normalized CCG, overlapping normalized CCG profile, peak time, physical distance between upstream and downstream synaptic pairs, converging and diverging properties. **d**, Similar plotting for putative inhibitory connections as of a central peak followed by an inhibitory trough. Note that these connections report a majority of inhibitory-to-inhibitory contact (92%).

**Supplementary Figure 6:**
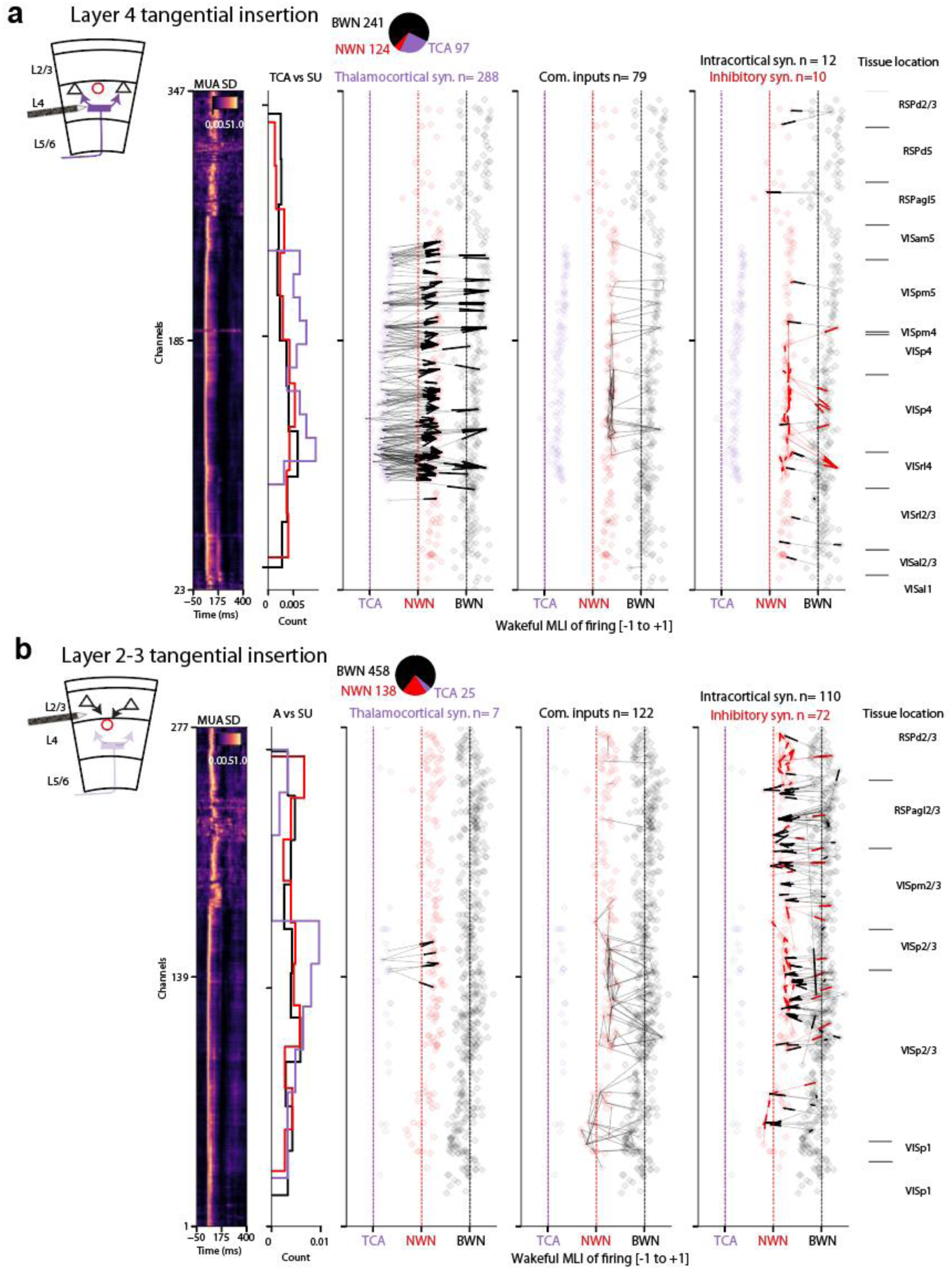
Stereotypical tangential insertions in layer 4 versus layer 2-3. **a**, Overview of a tangential recording within layer 4 of the cortex. The plots are made to overlay the physical location of the different observed CCG relationships. From left to right the plots are aligned to overlay several properties: Multi-unit-activity (MUA SD) showing which channels exhibit visual-driven activity (left). The location of TCA, NWN, and BWN numbers (histogram, left). The related excitatory monosynaptic pairs are illustrated by a thin line with a thickened end, illustrating the direction of the contacts (middle left and middle right). In each vertical connectivity plot TCA, NWN, or BWN are placed on their x-axis according to their modulation index of firing induced by wakefulness; the dotted line indicates the 0 modulation. TCA inputs are shown (middle left), aligned vertically with common inputs (middle) and intracortical excitatory monosynaptic contacts (middle right), together with the sharp-track tissue reconstruction (right). **b**, Similar plotting in a tangential recording within layer 2-3, highlighting the predominant presence of intracortical neurons toward layer 4 inhibitory neurons.

**Supplementary Figure 7:**
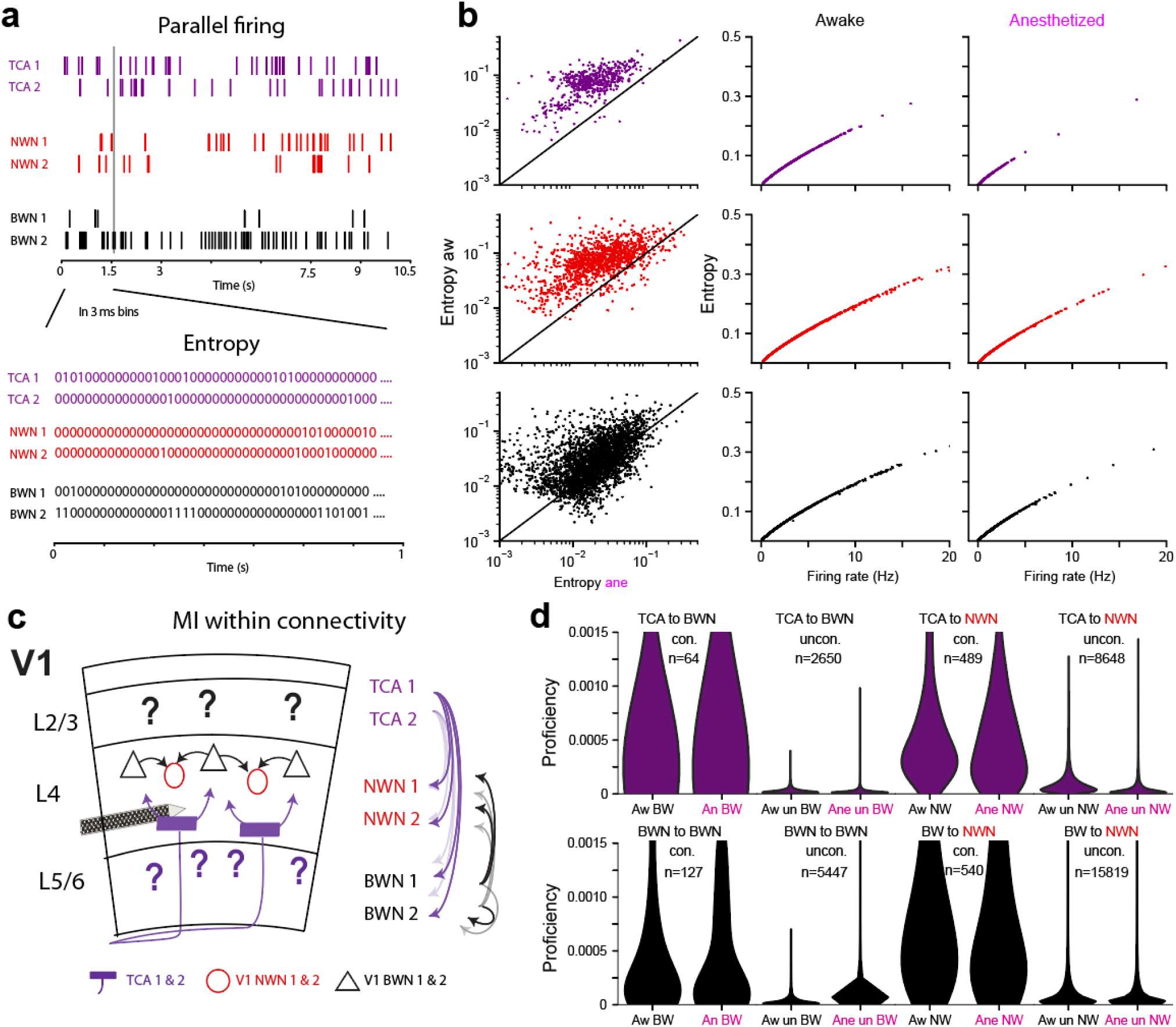
Entropy changes under anesthesia and proficiency within mono-synaptic pairs. **a**, Schematic of how to extract the entropy of each single individual neuron. **b**, Quantification of the entropy of single neurons in awake and anesthetized states (left columns, note the similarity of pattern with the awake/anesthetized firing changes in fig. 1). The entropy of each neuronal spike train has a strong relationship to its firing rate independent of wakeful (middle) or anesthetized states (right column). **c**, Schematic of excitatory mono-synaptic connected pairs used to calculate synaptic proficiency. **d**, Averages of proficiency within both types of excitatory synaptic connection (con., purple thalamocortical connections top line, black intracortical synapses bottom line), in awake versus anesthetized periods (pink labels), next to random selections of similar types of connections but to unconnected neuronal pairs (uncon.).

**Supplementary Figure 8:**
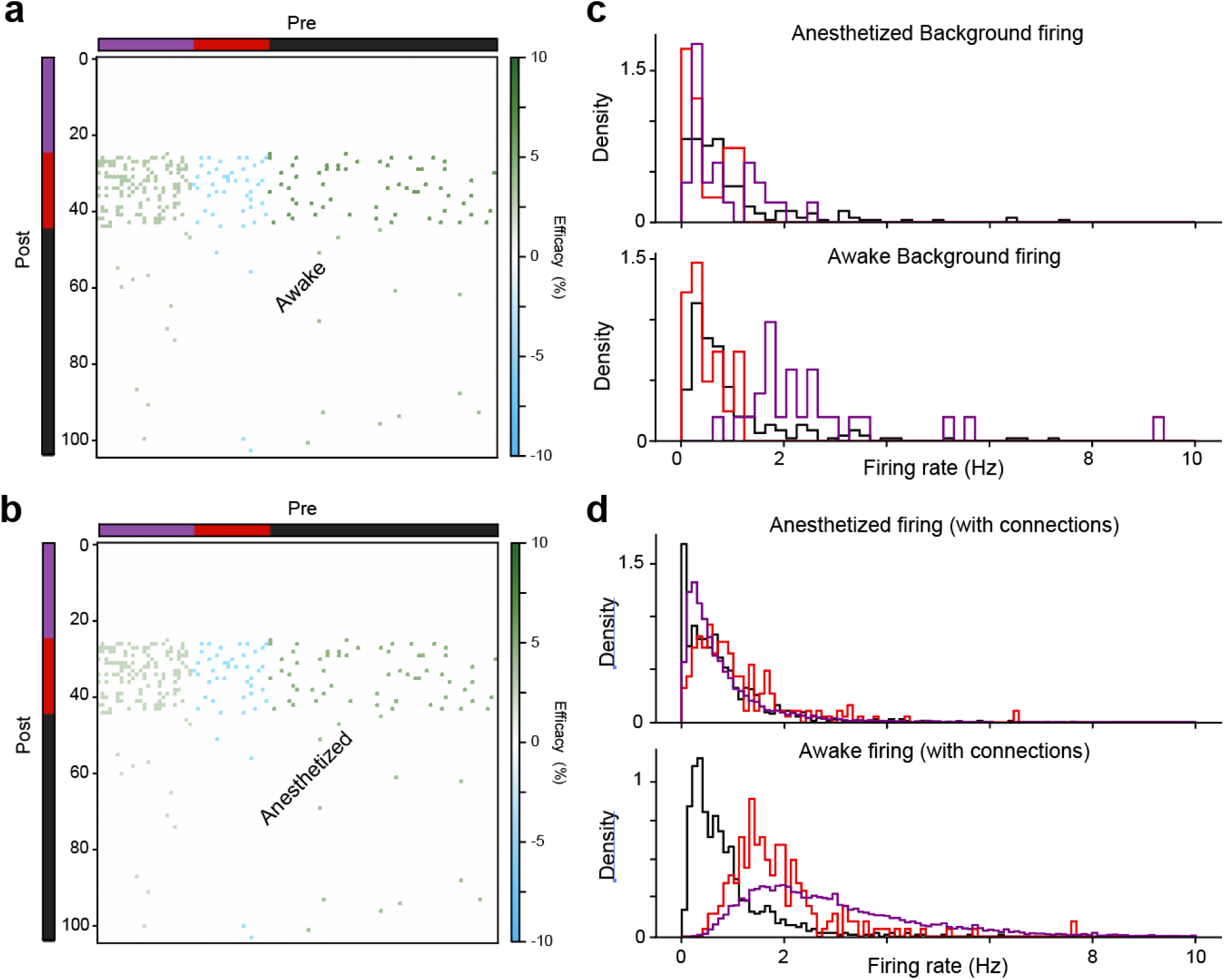
Modelling details, connectivity and wakeful modulations of firing levels. **a**, Connectivity matrix in awake state. **b**, Connectivity matrix in anesthetized state; note the slight color decrease corresponding to the slight decrease of efficacy under anesthesia in thalamocortical and intracortical connections. **c**, Background firing rate used in the model to account for unmonitored inputs (lognormal Poisson distribution). **d**, Firing rate obtained once the connections are established in anesthetized (top) and wakeful (bottom) contexts. Note the slight increase of firing under anesthesia, and the strong firing in wakeful conditions, which is mostly led by the higher TCA firing and its correlated activity.

**Supplementary Figure 9:**
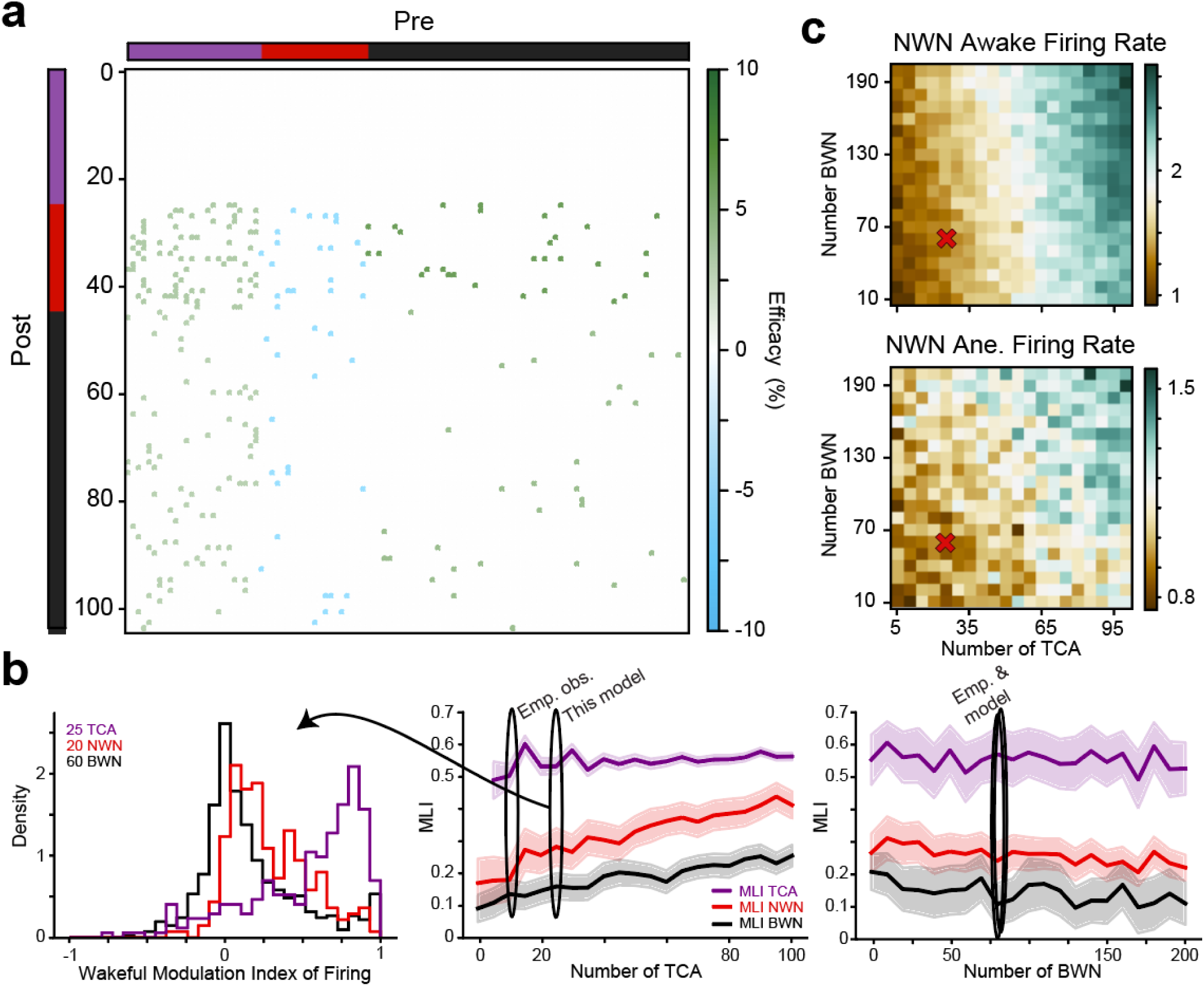
Modelling homogeneous BWN vs NWN connectivity. **a**, Connectivity matrix in awake states with similar numbers of connections to BWN and NWN. **b**, Wakeful MLI of firing of TCA, NWN and BWN having 25 TCA (left), MLI for different TCA numbers (middle), and different BWN numbers (right). Note the homogeneous increase of NWN MLI which does not reach 0.47 even with 110 TCA. **c**, Phase diagram of NWN wakeful firing for different BWN and TCA numbers with wakeful TCA firing level (top), and anesthetized (right).

